# *Astragalus* Saponins, Astragaloside VII and Newly Synthesized Derivatives, Induce Dendritic Cell Maturation and T Cell Activation Through IL-1β Production

**DOI:** 10.1101/2021.07.29.454313

**Authors:** Nilgun Yakubogullari, Ali Cagir, Erdal Bedir, Duygu Sag

## Abstract

Astragaloside VII (AST VII), a plant triterpenoid saponin isolated from *Astragalus* species, shows promise as vaccine adjuvant, as it supports a balanced Th1/Th2 immune response. However, the underlying mechanisms of its adjuvant activity have not been defined. Here we investigated the impact of AST VII and its newly synthesized semi-synthetic analogs on human whole blood cells, as well as on mouse bone marrow-derived dendritic cells (BMDCs). Cells were stimulated with AST VII and its derivatives in the presence or absence of LPS or PMA/ionomycin and the secretion of cytokines and the expression of activation markers were analyzed by ELISA and flow cytometry, respectively. AST VII and its analogs increased the production of IL-1β in PMA/ionomycin stimulated human whole blood cells. In LPS-treated mouse BMDCs, AST VII increased the production of IL-1β and IL-12, and the expression of MHC II, CD86, and CD80. The strength of the IL-1β boost correlated directly with the hydrophobicity of the AST VII compounds. In mixed leukocyte reaction, AST VII and derivatives increased the expression of the activation marker CD44 on mouse CD4^+^ and CD8^+^ T cells. In conclusion, AST VII and its derivatives strengthen pro-inflammatory responses, support dendritic cell maturation, and T cell activation *in vitro*. Our results provide insights into the mechanisms of the adjuvant activities of AST VII and its analogs, which will be instrumental to improve their utility as vaccine adjuvant.

## Introduction

Vaccination is one of the best strategies to induce protective immunity against infectious diseases and cancer. Although discovering a safe and effective antigen is the main target of vaccinology, the development of an adjuvant has also been gaining equal importance in recent years. Inactivated antigens, recombinant or subunit vaccines, containing purified parts of the antigens, are favored because of their safer profiles, but the poor immunogenic response requires utilization of an adjuvant (1).

QS-21, a triterpenoid saponin isolated from *Quillaja saponaria* stated as golden standard of saponin based adjuvants, has been in clinical trials as vaccine adjuvant for cancer, Alzheimer’s disease, malaria, HIV, etc. (2). Moreover, semi-synthesis and structure-activity relationship studies on QS-21 were carried out in order to overcome commercial development problems of QS-21, increase its efficacy in vaccine formulations, reduce its toxicity and reveal a rational path forward design of novel saponin adjuvants (2–6). GPI-0100, one of the promising semi-synthetic analogs of QS-21, was obtained by replacing the acyl group at the fucopyranosyl residue with dodecylamine linked glucuronic acid (at C6) located at C3 of the aglycone. Eleuterio et al., reported that GPI-0100 generated Th1 immunity and CTL response similar to QS-21 but with less toxicity (7).

Although several studies were undertaken to reveal the action mechanism of QS-21 in the last decade, the exact pathway was not established conclusively. In general, QS-21 was shown to intercalate with cell membrane cholesterols resulting in pore formation, acceleration of the antigen uptake by APCs, induction of cross-presentation through lysosomal destabilization, and production of lipid bodies to facilitate proteasomal route antigen degradation. Also, activation of NLPR3 inflammasome and further secretion of IL-1β and IL- 18 had a role in the cross-presentation induced by QS-21 (8–13).

*Astragalus* saponins have been widely used in Chinese Traditional Medicine for their pharmacological properties such as immunomodulatory, antioxidant, antitumor, and anti-viral properties (14). One of the cycloartane-type triterpenic saponins isolated from *Astragalus trojanus* (15), Astragaloside VII (AST VII), demonstrated a Th1/Th2 balanced immune response through antigen specific IgG, IgG1, and IgG2b antibody response, production of IL- 2, IFN-γ, TGF-β, IL-17A and enhanced splenocyte proliferation *in vitro* and *in vivo* studies (16–20). Besides inducing cellular and humoral immune response, high solubility in water, very low hemolytic activity at high concentrations, high stability, suitability to lyophilization make AST VII a potent adjuvant candidate. However, its mechanism of action has not been defined yet.

In this study, we prepared anologs of AST VII, namely DC-AST VII and DAC-AST VII, with the aim to find more potent ones and shed light on possible mechanisms contributing to their adjuvant actions. AST VII and its derivatives predominantly induced IL-1β production in human whole blood cells, dendritic cells and macrophages, and subsequently prompted dendritic cell maturation as well as T cell activation. The findings provided here are the first evidence in regards to the mechanism of action of AST VII and its analogs, and encourage further development of these compounds as potential vaccine adjuvant.

## Materials and Methods

### Reagents, cells and mice

AST VII was donated by Bionorm Natural Products, Izmir, Turkey. QS-21 was purchased from Desert King International (San Diego, CA, USA).

GM-CSF (Granulocyte Macrophage Colony Stimulating Factor) (R&D Systems, Minneapolis, USA), M-CSF (Macrophage Colony Stimulating Factor) (PeproTech, London, UK), LPS (Lipopolysaccharide) (invivogen, San Diego, CA, USA), Cytofix-fixation buffer (BD Bioscience, San Jose, CA,USA), Mouse IL-12p70, IL-2 ELISA kits (eBioscience), Human IL-2, IL-4, IL-17A, TNF-α, IFN-γ, IL-1β ELISA kits (eBioscience, San Diego, CA, USA), EasySep Mouse Naive CD4^+^ T cell and CD8^+^ T cell Isolation Kit (StemCell Technologies, Vancouer, Canada) were purchased.

Mouse bone marrow derived dendritic cells (BMDCs) were maintained in R5 medium containing RPMI 1640 medium with 5% Fetal Bovine Serum and 50 U/mL Penicillin/streptomycin (Gibco, Waltham, MA, USA) with 5 ng/mL GM-CSF. Mouse bone marrow derived macrophages (BMDMs) were cultured in R5 medium + L929 medium (1:1) supplemented with 10 ng/mL M-CSF.

C57BL/6 and BALB/c mice were housed in the animal facility of the Izmir Biomedicine and Genome Center (IBG). The mice were maintained in groups of 5 under standard conditions of temperature 22±1°C with regular 12 h light and 12 h dark cycles and free access to standard laboratory food and water. The experimental protocols were approved by the Local Ethics Review Committee for Animal Experimentation of IBG.

### Semi-synthesis studies of AST VII analogs and characterization of their physicochemical properties

New AST VII analogs were synthesized. Dicarboxylic AST VII (DC-AST VII) was synthesized through TEMPO mediated oxidation reaction of primary alcohols on glucose moieties to carboxylic acid (21). Dodecylamine conjugated AST VII (DAC-AST VII) was derived by conjugating free dodecylamines into glucuronic acid carboxyl of DC-AST VII via amide formation. Semi-synthetic compounds were extracted and further purified by column chromatography (Supplemental Table 1). The chemical structures of the analogs (Fig. 1) were elucidated by 1D and 2D NMR [Varian MERCURY plus AS400 (400 MHz)] and HRTOF-MS spectroscopy (Agilent 1200/6530) (Supplemental Table 1).

**FIGURE 1.**
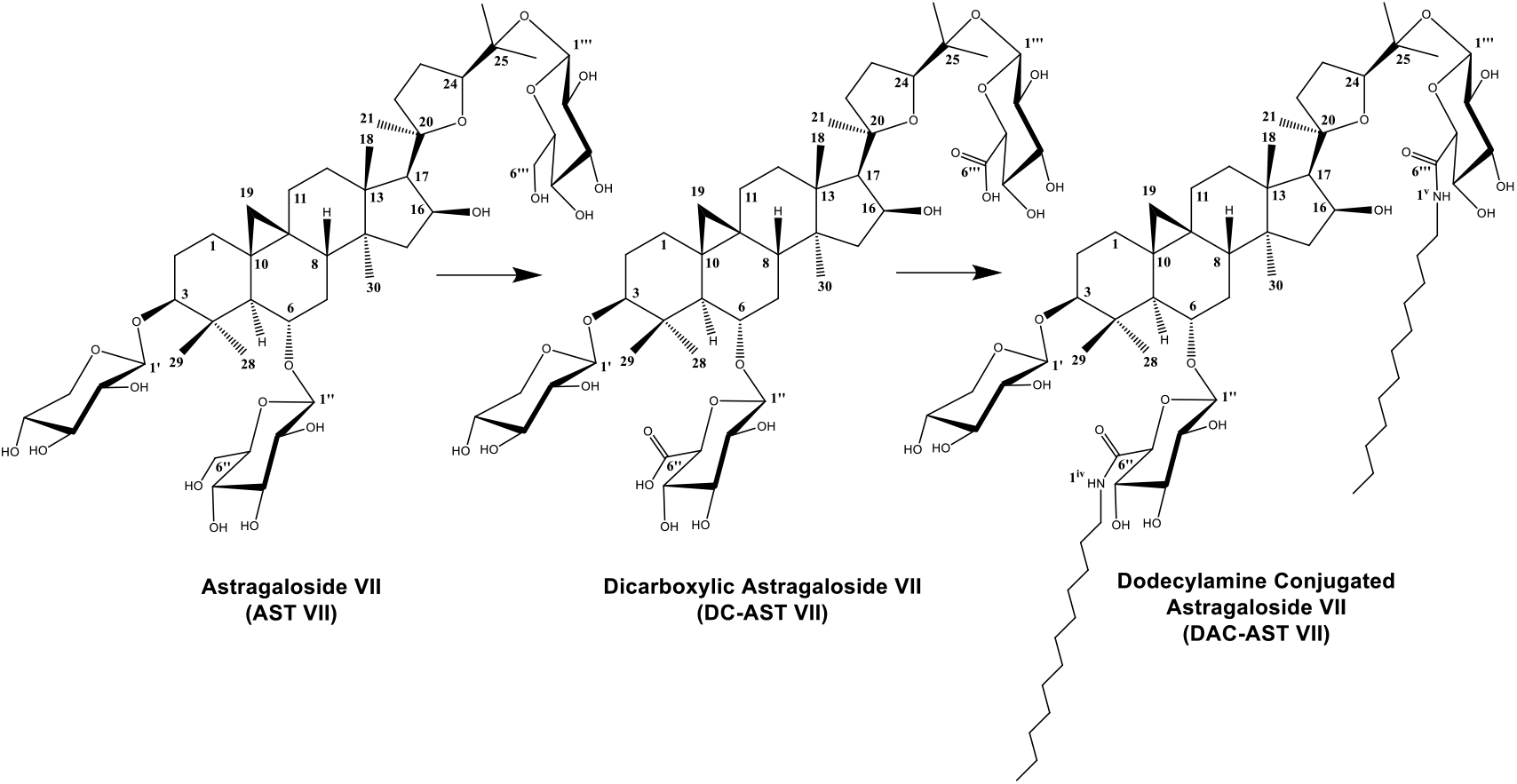
Chemical structures of AST VII and newly synthesized derivatives: Dicarboxylic Astragaloside VII (DC-AST VII) and Dodecylamine Conjugated Astragaloside VII (DAC-AST VII) To determine critical micelle formation, 1 mg/mL dansyl chloride dissolved in acetone (20 μL) was transferred to 96 well plates and left to evaporate. A series of concentrations of AST VII and its derivatives (0.01 μM to 20 mM) were added into solvent evaporated dansyl chloride and incubated overnight by shaking in the dark and at room temperature. The fluorescent intensity was measured by a microplate reader at excitation and emission wavelengths filter of 335 nm and 518 nm, respectively (Thermo, Varioskan Flash).

To assess the particle size distribution and zeta potential of the micelles, dynamic light scattering measurement was performed using a Zetasizer Nano ZS at a 173° scattering angle at 25°C. AST VII (4000 μM), DC-AST VII (10 μM) and DAC-AST VII (10 μM) were dissolved in water, and the particle sizes were shown as number of particle distribution.

### Cytotoxicity of AST VII analogs

Human endometrial carcinoma (HeLa), human lung adenocarcinoma (A549), human prostate carcinoma (Du145), human ductal carcinoma (HCC-1937), human breast adenocarcinoma (MCF-7), and human lung fibroblasts (MRC-5) were obtained from the American Type Culture Collection and maintained as exponentially growing monolayers by culturing according to the supplier’s instructions. For the cytotoxicity analysis, each cell line was exposed to the compounds (DC-AST VII and DAC-AST VII) at a final concentration of 2, 4, 8, 16, or 32 μM for 48 h. After 48 h incubation, the cell viability was analyzed by the MTT assay (Sigma Aldrich, St. Louis, Missouri, ABD) according to the manufacturer’s instructions. DMSO was used as a negative control. Absorbance was measured with a microplate reader at 570 nm (Varioscan, Thermo Fisher Scientific, US). IC_50_ values, defined as the compound concentration at which the response level was reduced to half its maximum, were calculated using GraphPad Prism 6 (San Diego, CA, USA).

### Hemolytic activity

Hemolytic activity assay was performed with human red blood cells obtained from healthy volunteers (16). 7 mL of blood was washed and centrifuged three times at 2000×g for 5 min with sterile PBS. The cell suspension was diluted in saline solution with a final concentration of 0.5%. 0.01 mL of the cell suspension was mixed separately with 0.01 mL saline solution of AST VII and derivatives at varying concentrations (0, 25, 50, or 250 μg/mL). The mixtures were incubated for 30 min at 37°C and centrifuged at 800×g for 10 min. The free hemoglobin content of the supernatants was measured spectrophotometrically at 540 nm. Saline and distilled water were referred to minimum and maximum hemolytic controls.

### Human whole blood (hWB) stimulation assay

Human whole blood (hWB) stimulation assay was performed to demonstrate the cytokine release profiles of AST VII analogs in parallel to AST VII and QS-21. Heparinized whole blood from healthy volunteers was supplemented in 1:10 and 1:20 with RPMI-1640 medium, 100 U/mL penicillin/streptomycin and 10% fetal bovine serum. PMA (phorbol 12-myristate 13-acetate) (50 ng/mL) and ionomycin (400 ng/mL) (Sigma, St. Louis, Missouri, USA) stimulated hWB were treated with the following compounds AST-VII, DC-AST VII, DAC-AST VII and QS-21 at the concentrations of 2, 4, 8, 16, 32 μg/mL. The cells were incubated at 37°C in 5% CO_2_ for 48 h. Supernatants of each well were collected and stored at − 20°C for ELISA.

### Generation of bone marrow derived dendritic cells (BMDCs) and macrophages (BMDMs)

BMDCs were prepared as described previously (22). Briefly, femurs and tibias of C57BL/6 and BALB/c mice were collected and flushed with sterile DPBS twice. The resulting bone morrow cells were resuspended in R5 medium (RPMI 1640 medium containing 5% heat-inactivated FBS, 2 mM L-glutamine, 50 U/mL penicillin/streptomycin) plus 5 ng/mL GM-CSF. A total of 2×10^6^ cells per bacteriological culture plate were cultured for 10 days. The cells were transferred to a new plate on day 6 and fresh medium was added to the culture on days 3 and 8. Non-adherent and loosely adherent cells were collected on day 10 and the cells were verified to be 80% CD11c^+^ by flow cytometry analysis. BMDMs were prepared as described previously (23). Briefly, bone marrow cells from C57BL/6 mice were resuspended in R5 medium containing 10 ng/mL M-CSF and cultured for a day in a tissue culture plate. The next day, the non-adherent cells were collected and transferred to a 6-well low attachment plate in 4 mL R5 medium containing 30% L929 conditioned medium and 10 ng/mL M-CSF per well for 7 days. Fresh medium was added to the culture on day 6. After that, the cells were purified by centrifugation over Ficoll-Paque plus. The cells were verified to be >90% CD11b^+^ F4/80^+^ by flow cytometry analysis.

### Stimulation of BMDCs and BMDMs with AST VII, DC-AST VII, and DAC-AST VII

BMDCs were suspended in R5 medium and cultured in 96 well plate at the concentration of 2.5×10^5^ BMDCs/well in 200 μL. AST VII and its derivatives were dissolved in DMSO. Different concentrations of AST VII (1 to 64 μM), DC-AST VII and DAC-AST VII (2 to 20 μM) were added to the appropriate wells and incubated for 24 h at 37°C in 5% CO_2_ incubator. In the presence of LPS (10 ng/mL), BMDCs were treated with AST VII, DC-AST VII and DAC-AST VII (0.5, 2, 2.5, 5, 10 μM) and incubated for 24 h. The cells were analyzed for the expression levels of MHC II, CD86 and CD80 by flow cytometry and the supernatant was analyzed for IL-12 production by ELISA. BMDMs were plated as 1×10^5^ BMDMs/well in 200 μL R5 medium. The cells were treated with LPS (10 ng/mL) plus different concentrations of AST VII, DC-AST VII and DAC-AST VII (2, 5, 10 μM). 6 hours later, the supernatant was collected and the IL-1β cytokine release was analyzed by ELISA.

### Naive CD4^+^ and CD8^+^ T cell isolation

Spleens from C57BL/6 mice were disrupted in DPBS containing 2% FBS. Single splenocyte suspensions were obtained by passing the cells through a 70 μm mesh nylon strainer, followed by centrifugation at 300xg for 10 min three times. The cell pellet was resuspended in DPBS containing 2% FBS with a final concentration of 1×10^8^ nucleated cells/mL. Naive CD4^+^and CD8^+^ T cells were isolated by negative selection using StemCell EasySep™ Mouse Naive CD4^+^ and CD8^+^ T Cell Isolation Kit following manufacturer’s instructions.

### Mixed lymphocyte reaction (MLR)

BMDCs generated from BALB/c mice were plated in 96 well plates as 2×10^4^ cells/well in 100 μL R5 medium. The cells were treated with LPS (10 ng/mL), AST VII (5 μM), DC-AST VII (5 and 10 μM), DAC-AST VII (5 and 10 μM) and incubated for 24 h at 37°C in 5% CO_2_ incubator. The next day, followed by washing with DPBS, naive CD4^+^ and CD8^+^ T cells isolated from C57BL/6 were added to the 96 well plates in 1:5 ratio (BMDCs: T cells). The cells were incubated for 3 days at 37°C in 5% CO_2_ incubator. CD4^+^ and CD8^+^ T cell activation was assessed by expression of CD44 by flow cytometry. The cell supernatant was collected and stored at −20°C for subsequent cytokine analysis.

### Flow cytometry

For the staining of cell surface molecules, cells were resuspended in FACS buffer (PBS, 1% BSA, 0.025% NaN3) and Fcγ receptors were blocked with 1/200 diluted anti-mouse CD16/32 antibodies (Fc block) (Tonbo Biosciences, San Diego, CA, USA) for 10 min on ice. BMDCs and splenocyte single cell suspensions were stained with the following 1/400 diluted fluorochrome conjugated anti-mouse Abs: anti-CD11c-APC, anti-MHC (I-A/I-E)-PE-Cy7, anti-MHC (I-A/I-E)-APC-Fire750, anti-CD80-Pacific Blue, anti-CD86- PerCp Cy5.5, anti- CD86-PE-Cy7, anti-F4/80-PE, anti-CD11b-PerCp-Cy5.5 (Biolegend, San Diego, CA, USA) and incubated in the dark on ice for 40 minutes. The cells were fixed with Cytofix-fixation buffer (BD Biosciences) for 15 min on ice and cell fluorescence was assed using Fortessa (BD Biosciences). The data were analyzed with FlowJo software version 10. Gating strategy for BMDCs: FSC-A: SSC-A > singlets > CD11c^+^ MHC II^+^ > MHC II^+^ CD86^+^, MHC II^+^ CD80^+^. Gating strategy for DCs from splenocytes: FSC-A: SSC-A > singlets > F4/80^−^ > CD11b^+^ CD11c^+^ > MHC II^+^ CD86^+^ > MHC II^+^ CD80^+^. Gating strategy for lymphocytes: FSC-A: SSC-A > singlets > live-dead > CD4^+^ CD8^−^; CD4^−^CD8^+^ > CD44^+^.

### ELISA

The cell supernatants were collected and the concentration of human IL-2, IFN-γ, IL-4, IL-17A, TNF-α, IL-1β and of mouse IL-12 and IL-1β were assessed by ELISA (eBioscience, San Diego, CA, USA) in accordance with the manufacturer’s instructions.

### Statistical analysis

Data for all experiments were analyzed with Prism software (GraphPad). The data are expressed as mean ± SD. Unpaired Student’s t-test, One-way ANOVA and Dunnett’s or Tukey’s multiple comparisons tests were used for comparison of experimental groups when appropriate. P values of less than 0.05 were considered statistically significant.

## Results

### Cytotoxic and hemolytic activities of AST VII analogs on human cancer cell lines and erythrocytes

The cytotoxic and hemolytic activities of the newly synthesized AST VII analogs were investigated. The lead compound, AST VII, did not reveal any cytotoxic or hemolytic activities in previous studies (16). Five cancer cell lines (A549, Du145, HCC-1937, HeLa, MCF-7) and one control cell line (MRC-5) were used to screen the cytotoxic properties of DC-AST VII and DAC-AST VII by MTT assay. DC-AST VII slightly reduced the cell viability compared to DMSO control in cancerous cell lines except HCC-1937, which had more prominent reduction at the concentrations of 8 and 32 μM. Moreover, DC-AST VII at 2 μM concentration statistically enhanced cell proliferation compared to DMSO control in MRC-5 cells (*p* < 0.001) In contrast to DC-AST VII, DAC-AST VII demonstrated cytotoxicity in all cell lines in a dose-dependent manner with the following IC50 values: 29.51 μM for A549, 7.91 μM for Du145, 12.06 μM for HCC-1937, 11.6 μM for HeLa, 9.79 μM for MCF-7 and 17.04 μM for MRC-5 (Supplemental Fig. 1).

As one of the specific properties of saponins is hemolysis, next the hemolytic activity of AST VII analogs on human erythrocytes was investigated. Serially diluted (1/5) concentrations (250 μM to 0.4 μM) of DC-AST VII and DAC-AST VII were analyzed. Only DAC-AST VII at 50 μM and DC-AST VII at 10 μM demonstrated slight hemolytic activity compared to the control saline treatment (*p* < 0.01 and *p* < 0.05) (Supplemental Table 2). As known from previous studies, AST VII at the concentration of 2.5 to 500 μg/mL does not cause hemolysis of rabbit red blood cells (19). Overall, our data suggest that the structural alteration at the C-6 position of the glucose units of AST VII caused a minor difference in its hemolytic activity and that the hydrophobicity of the compounds and presence of the fatty acid chains are important structural features defining the cytotoxic properties of AST VII analogs.

### Evaluation of immunomodulatory activities of AST VII and its derivatives based on cytokine release on human whole blood

The human whole blood (hWB) stimulation assay is a simple and effective approach to evaluate the immunomodulatory activities of the test compounds as it contains diverse immune cells, like T cells, B cells, NK cells, monocytes and granulocytes (24). To address what type of cellular immune response (Th1/Th2/Th17) was induced by AST VII and its analogs, the cytokine release profiles (IL-2, IFN-γ, IL-17A, IL-1β, TNF-α, IL-4) of PMA-ionomycin (P+I) stimulated hWB were investigated. As PMA (protein kinase C activator) is commonly used with ionomycin (calcium ionophore) to activate T cells by inducing NF-κB and NFAT transcription factors and, successively leading to production of cytokines (25), hWB was stimulated with P+I.

hWB was diluted in 1/10 ratio with RPMI-1640 medium and stimulated with PMA (50 ng/mL) and ionomycin (400 ng/mL) in the presence or absence of AST VII or its derivatives. QS-21, a widely used saponin based adjuvant was also tested for comparison. Surprisingly, in the absence of the saponin compounds we did not observe any significant production of IL-1β and IL-17A cytokines compared to control group (untreated cells) (Fig. 2A, 2B). This was thought to be caused by a too high cell concentration in the 1/10 diluted hWB. Therefore, for later analysis we decided to dilute hWB in 1/20 ratio. When the results are analyzed in detail, AST VII (up to 2.24 fold), DC-AST VII (2.52 fold), and DAC-AST VII (3.32 fold) substantially increased IL-1β production (Fig. 2A). Overall, the increase in IL-1β production induced by AST-VII and its derivatives was higher compared to QS-21 (Fig. 2A). In terms of IL-17A production, AST VII (up to 5.0o5 fold) and QS-21 increased the production of IL-17A compared to P+I treatment alone (*p* < 0.05, *p* < 0.01, *p* < 0.001) (Fig. 2B).

**FIGURE 2.**
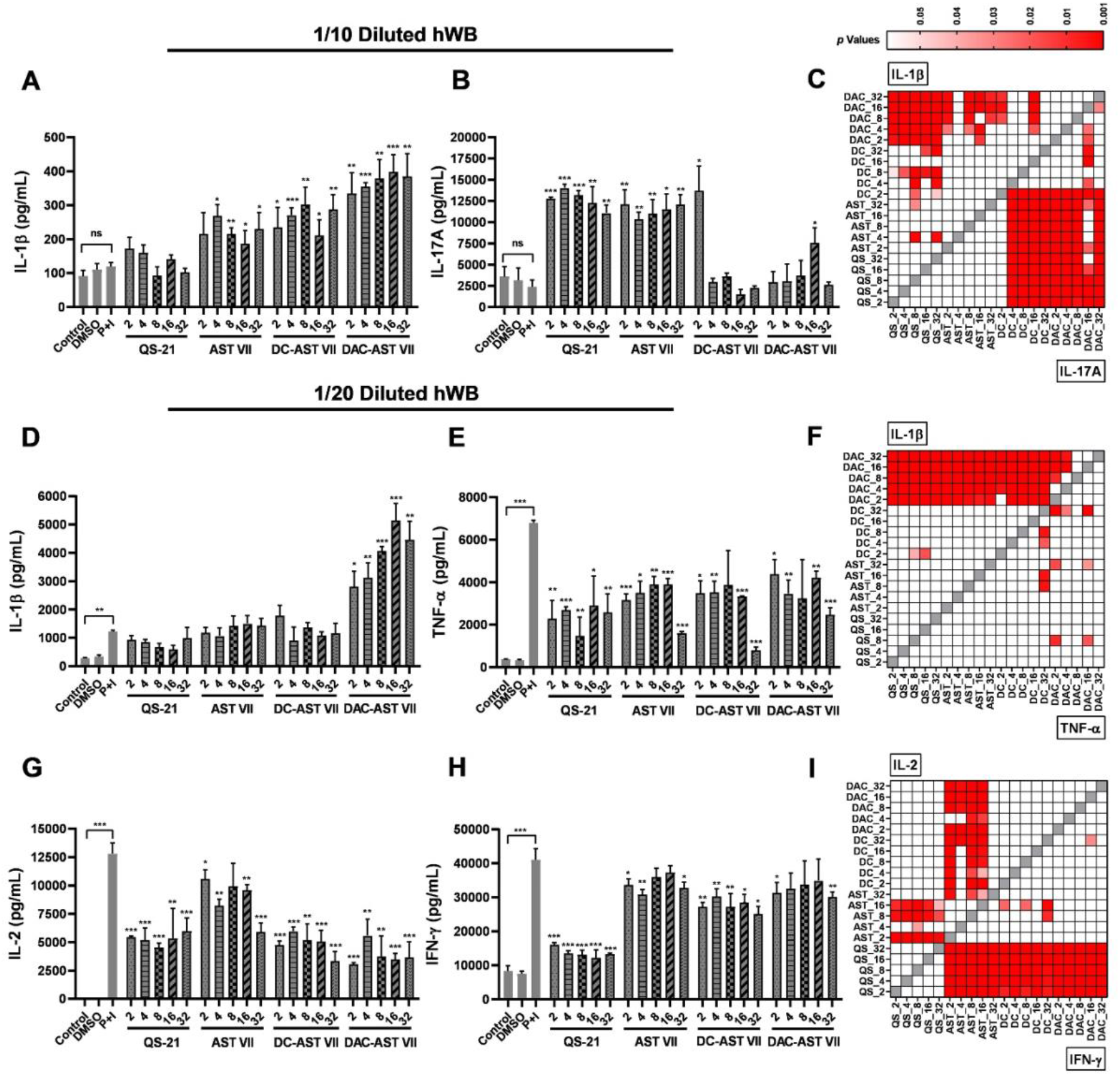
AST VII and its derivatives (DC-AST VII and DAC-AST VII) alter the production of pro-inflammatory cytokines by hWB cells. Diluted hWB was co-treated with PMA (50 ng/mL) and ionomycin (400 ng/mL) and one of the following compounds: QS-21, AST VII, DC-AST VII, DAC-AST VII, at the concentration of 2-32 μg/mL for 48 h. The supernatants were collected for the detection of cytokines by ELISA. **(A)** IL-1β in 1/10 diluted hWB, **(B)** IL-17A in 1/10 diluted hWB, **(C)** Heat map of IL-1β and IL-17A represents the *p* values calculated by analyzing different compounds with each other or analyzing each compound with respect to their different concentrations (2-32 μg/mL). **(D)** IL-1β in 1/20 diluted hWB, **(E)** TNF-α in 1/20 diluted hWB, **(F)** Heat map of IL-1β and TNF-α represents the *p* values in 1/20 diluted hWB, **(G)** IL-2 in 1/20 diluted hWB, **(H)** IFN-γ in 1/20 diluted hWB, **(I)** Heat map of IL-2 and IFN-γ represents the *p* values in 1/20 diluted hWB. DMSO was used as vehicle control. Data shown are mean ± SD of triplicates and representative of two independent experiments with similar results. Statistical analyses were performed between treated groups and P+I (PMA+Ionomycin) by One-way ANOVA and Student-t test and, in treated groups between each other for each concentration by Tukey’s multiple comparison tests. \**p* < 0.05, \*\**p* < 0.01, \*\*\**p* < 0.001, not statistically significant (ns). DAC: DAC-AST VII, DC: DC-AST VII, AST: AST VII, QS: QS-21.

The correlation between treated compounds and their concentration on cytokine production was demonstrated as a heat map of *p* values. Hence, DAC-AST VII induced more IL-β than QS-21 at all concentrations (*p* < 0.001), and also AST VII and DC-AST VII at the concentration of 16 μM (*p* < 0.001). Moreover, QS-21 and AST VII treatments at 4 to 32 μM statistically augmented IL-17A levels compared to DC-AST VII and DAC-AST VII (*p* < 0.01, *p* < 0.001) (Fig. 2C). Also, DC-AST VII (2 μM) and DAC-AST VII (16 μM) were able to induce more IL-17A production (*p* < 0.05) compared to the P+I treated group, indicating that the production was also dependent on the compound concentration. IL-2, IFN-γ, TNF-α, and IL-4 cytokines could not be detected by ELISA.

In contrast to the 1/10 diluted hWB, the 1/20 diluted hWB demonstrated statistically significant cytokine production (IL-1β, TNF-α, IL-2, IFN-γ) after P+I treatment, compared to the control group (untreated cells) (*p* < 0.01, *p* < 0.001). AST VII and DC-AST VII did not enhance IL-1β production, whereas DAC-AST VII (at 16 μg/mL, 4.2 fold) remarkably boosted IL-1β levels compared to P+I alone (*p* < 0.001) (Fig. 2D). The production of TNF-α, IFN-γ, and IL-2 cytokines was suppressed following the treatment with AST VII, DC-AST VII, or DAC-AST VII. There was no consistency in the TNF-α production with respect to the concentration of compounds (Fig. 2E). DAC-AST VII was more effective than the other compounds to induce IL-1β production (*p* < 0.001). Moreover, DAC-AST VII at the concentration of 16 and 32 μg/mL demonstrated significant increase compared to DAC-AST VII at 2 and 4 μg/mL (*p* < 0.001 and *p* < 0.01) (Fig. 2F), demonstrating a dose-dependent increase in IL-1β. For the IL-2 response, AST VII was superior compared to the other compounds (Fig. 2G). No difference between AST VII and its analogs was observed for IFN-γ production (Fig. 2H, Fig. 2I). The heat map for IL-2 shows that AST VII at the concentrations of 2, 8, and 16 μg/mL was superior in IL-2 induction compared to QS-21, DC-AST VII, and DAC-AST VII (*p* < 0.001). AST VII at 2, 8, and 16 μg/mL also induced higher IL-2 levels than 32 μg/mL (p < 0.001, p < 0.01, p < 0.05, respectively) (Fig. 2I), showing that low concentrations of AST VII were more effective for IL-2 induction than higher concentrations. AST VII, DC- AST VII, and DAC-AST VII stimulated IFN-γ production more effectively than QS-21 (*p* < 0.001) (Fig. 2I). Low levels of IL-17A was detected, but the levels of IL-4 could not be measured by ELISA. These data show that when co-treated with PMA-ionomycin, AST VII and derivatives altered the cytokine profile of hWB. Overall, AST VII induced the production of the pro-inflammatory cytokines such as IL-1β and IL-17A, while its derivatives induced the production of only IL-1β.

### IL-1β secretion following treatment of AST VII and its derivatives in BMDCs and BMDMs

Next, we investigated the impact of AST VII and its derivatives on the production of IL-1β by innate immune cells, such as dendritic cells and macrophages (26). Bone marrow cells from C57BL/6 mice were differentiated into bone marrow-derived dendritic cells (BMDCs) or bone marrow-derived macrophages (BMDMs) *in vitro* in the presence of GM-CSF or M-CSF, respectively. LPS alone has been reported to be a weak stimulus for IL-1β secretion in dendritic cells (27). Upon co-treatment with LPS (10 ng/mL), AST VII and its derivatives significantly boosted IL-1β secretion in BMDCs compared to LPS alone (*p* < 0.01, *p* < 0.001). When the effect of the compound concentration on IL-1β production was investigated, only DAC-AST VII at the concentration of 2.5 and 10 μM made a statistically significant difference (*p* < 0.05). Moreover, the effect of the compounds on the production of IL-1β was examined at the constant concentration. AST VII (0.5 μM) induced higher IL-1β levels than DC-AST VII at 0.5 μM (*p* < 0.01), but not at 0.5 μM (ns). There was no statistically significant difference between AST VII and its derivatives at the concentration of 2.5 μM. AST VII (10 μM) substantially augmented IL-1β production compared to DC-AST VII (10 μM) and DAC-AST VII (10 μM) (*p* < 0.001) (Fig. 3A).

**FIGURE 3.**
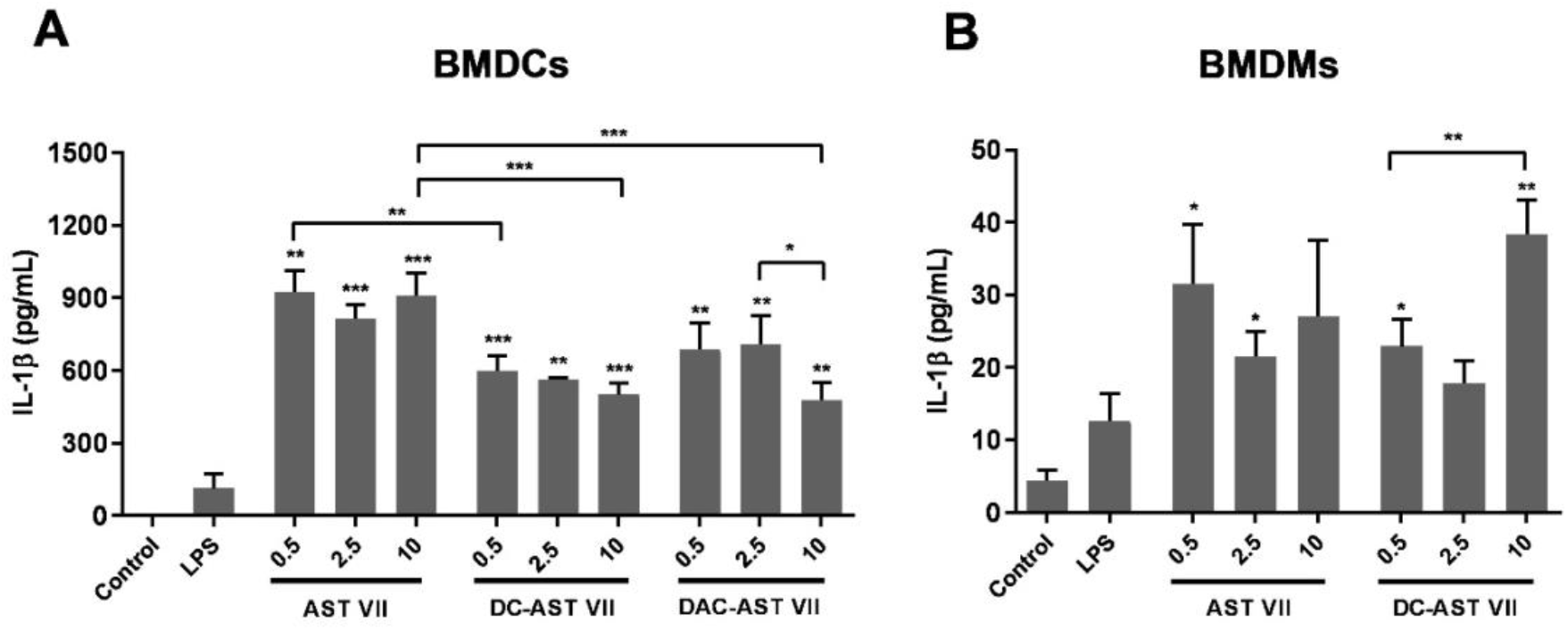
AST VII and its derivatives induce IL-1β secretion in BMDCs and BMDMs. **(A)** BMDCs and **(B)** BMDMs generated from bone marrow cells of C57BL/6 mice were treated with LPS (10 ng/mL) alone or LPS with AST VII, DC-AST VII, or DAC-AST VII at the concentrations of 0.5, 2.5, or 10 μM for 6 h. Unstimulated cells were used as control. The cell culture supernatants were collected to analyze IL-1β concentrations by ELISA. Data shown are the mean ± SD of triplicates and representative of two independent experiments with similar results. Statistically significant differences were analyzed between treated groups and LPS by One-way ANOVA and Student-t test and, in treated groups between each other by Tukey’s multiple comparisons test. *p < 0.05, **p < 0.01, ***p < 0.001.

In BMDMs, AST VII and DC-AST VII demonstrated slight increase in IL-1β compared to LPS alone (*p* < 0.05; *p* < 0.01), whereas IL-1β production after DAC-AST VII treatment was below the detection limit (data not shown). 10 μM DC-AST VII induced significantly higher levels of IL-1βcompared to 0.5 μM (*p* < 0.01) (Fig. 3B). These data indicate that for BMDCs all compounds and for BMDMs AST VII and DC-AST VII boosted the production of IL-1β in the presence of LPS.

### AST VII and its derivatives induced dendritic cell maturation and activation

IL-1β is produced by activated dendritic cells and can lead to the activation of other dendritic cells (28). Given the enhanced production of IL-1β in hWB, BMDCs, and BMDMs by AST VII and its derivatives, we next investigated if these compounds affect dendritic cell maturation and activation. The maturation state of BMDCs was evaluated by the analysis of MHC II, CD86, and CD80 expression by flow cytometry and the secretion of IL-12 by ELISA.

The treatment of AST VII induced neither the expression of MHC II, CD86, or CD80 nor the production of IL-12 in BMDCs even at high concentrations (up to 60 μg/mL) (Supplemental Fig. 2A, 2B, 2C). To investigate whether AST VII enhances dendritic cell maturation/activation by the help of other cell types or secreted cytokines in the microenvironment, single cell suspension of splenocytes from C57BL/6 mice was treated with AST VII alone at concentrations of 5, 10, or 20 μg/mL. However, AST VII stimulation did not result in the upregulation of MHC II, CD80, or CD86 (Supplemental Fig. 2D). These findings demonstrate that AST VII alone does not affect maturation/activation of dendritic cells *in vitro*.

Fig. 3A shows that AST VII was able to significantly increase IL-1β production by BMDCs in the presence of the TLR4 agonist LPS (10 ng/mL). Therefore, BMDCs were incubated with LPS (10 ng/mL) with or without AST VII at the concentrations of 2, 5 or 10 μM for 24 h. Compared to LPS alone, AST VII plus LPS significantly increased the expression on BMDCs of CD86 (all doses, Fig. 4A), CD80 (2 μM, 10 μM, Fig. 4B), and MHC II (10 μM, Fig. 4C). AST VII at 10 μM was more effective than 2 μM or 5 μM in the upregulation of both CD86 and CD80 (*p* < 0.05 and *p* < 0.01, respectively) (Fig. 4A, B). Besides the upregulation of the costimulatory molecules, secretion of IL-12 is also one of the indicators of DC activation (29). The production of IL-12 by BMDCs was significantly increased following co-stimulation with LPS and AST VII compared to LPS alone (*p* < 0.01) (Fig. 4D). These data show that AST VII induces dendritic cell maturation and activation in cooperation with LPS (Fig. 4E).

**FIGURE 4.**
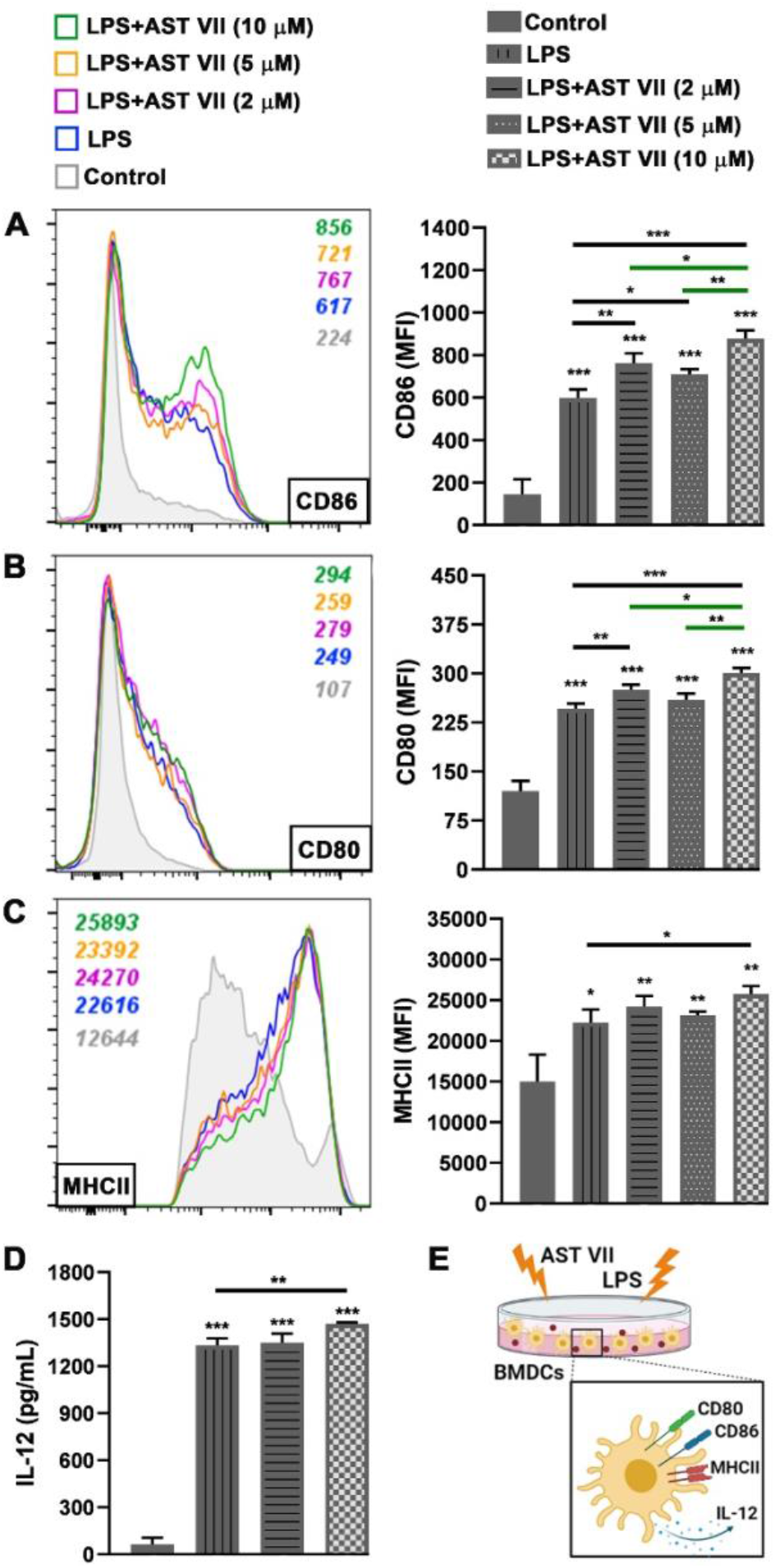
AST VII induces the maturation and activation of BMDCs in the presence of LPS. BMDCs generated from C57BL/6 mice bone-marrow cells were co-treated with LPS (10 ng/mL) and AST VII at the concentrations of 2, 5, or 10 μM for 24 h. The expression of **(A)** CD86, **(B)** CD80, and **(C)** MHCII on CD11c^+^ MHCII^+^ BMDCs was determined by flow cytometry. For A, B, and C, representative histogram (left panel) and summary graphs (right panel) are shown. **(D)** IL-12 titers were measured by ELISA. **(E)** Schematic illustration of DC maturation/activation after the treatment with AST VII and LPS. Data shown are mean ± SD of triplicates and representative of three independent experiments with similar results. Statistically significant differences of treated groups versus control (untreated cells) or LPS and, in treated groups between each other by One-way ANOVA and Student-t test are indicated. *p < 0.05, **p ≤ 0.01, ***p ≤ 0.001. *Created with* BioRender.com

To evaluate the impact of DC-AST VII and DAC-AST VII on dendritic cell maturation/activation, BMDCs were treated with the compounds for 24 h in the absence/presence of LPS. Without LPS stimulation, DC-AST VII at the concentration of 5 and 10 μM increased the expression of CD86, CD80, and MHC II compared to the control group (untreated) (*p* < 0.05) (Fig. 5A, 5B, 5C). In general, DAC-AST VII treatment alone did not demonstrate an increase in the expression of CD86, CD80, and MHC II (n.s.). Interestingly, MHC II expression on BMDCs was decreased by DAC-AST VII at the concentration of 10 μM compared to the control group (*p* < 0.05) (Fig. 5C).

**FIGURE 5.**
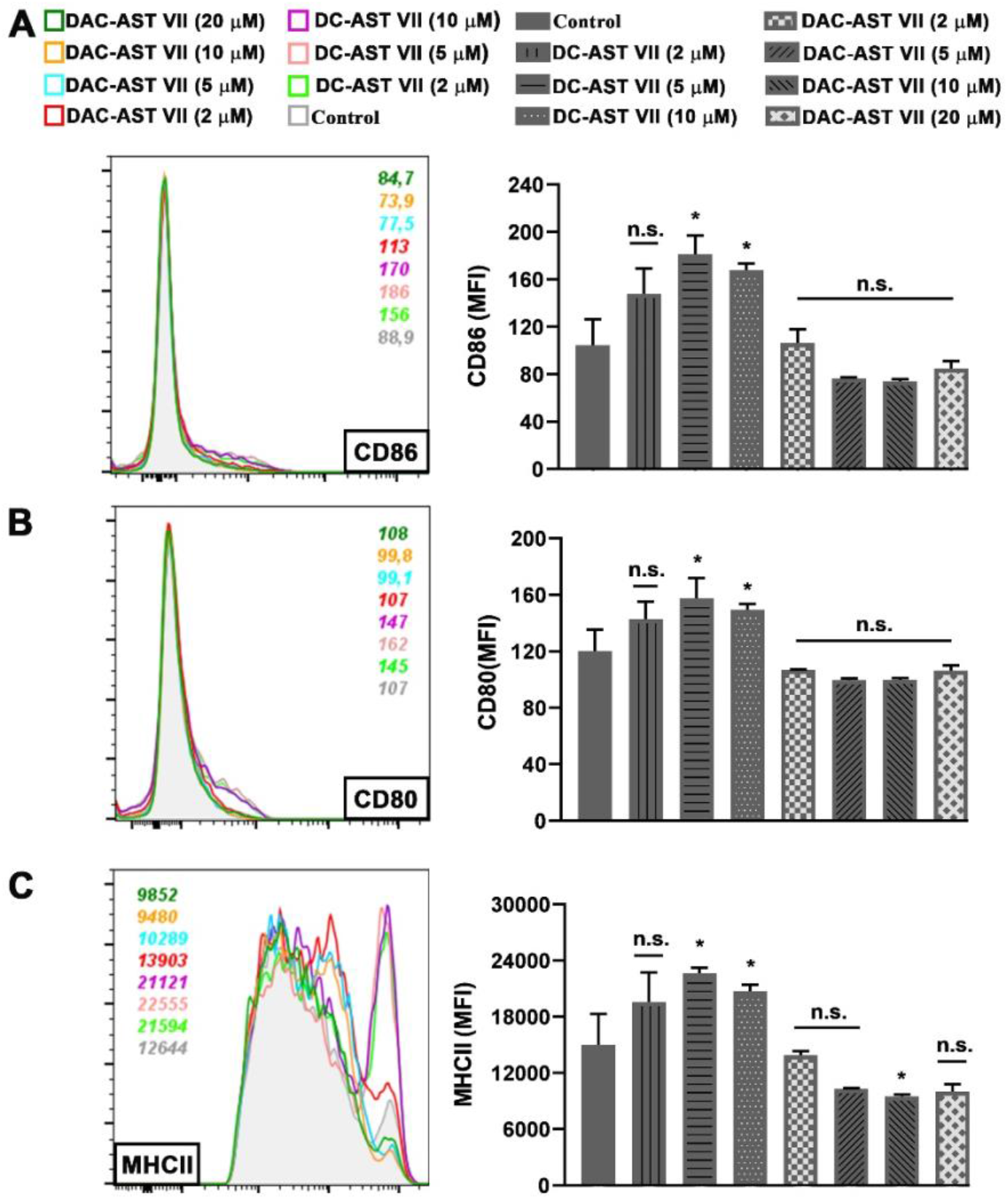
DC-AST VII but not DAC-AST VII slightly induce the maturation and activation of BMDCs in the absence of LPS. BMDCs generated from C57BL/6 mice bone marrow cells were treated with DC-AST VII or DAC-AST VII at the concentrations of 2, 5, 10, or 20 μM for 24 h. The expression of **(A)** CD86, **(B)** CD80, and **(C)** MHC II on CD11c^+^ MHCII^+^ BMDCs was determined by flow cytometry. For A, B, and C, representative histogram (left panel) and summary graphs (right panel) are shown. Data shown are mean SD of triplicates and representative of two independent experiments with similar results. Statistically significant differences of treated groups versus control (untreated cells) by One-way ANOVA and Student-t test, in treated groups between each other by Tukey’s multiple comparisons test are indicated. *p < 0.05, **p ≤ 0.01, ***p ≤ 0.001, n.s. (not statistically significant).

Next, to determine whether stimulation of DC-AST VII and DAC-AST VII with LPS would alter their adjuvant activity on BMDCs, the cells were co-treated with DC-AST VII or DAC-AST VII at 10 μM along with LPS (10 ng/mL). In the presence of LPS, DC-ASTVII and DAC-AST VII reduced the expression of CD86, CD80 and MHC II compared to LPS alone (*p* < 0.05, *p* ≤ 0.01, *p* ≤ 0.001). The reduction by DAC-AST VII was more prominent compared to DC-AST VII (Fig. 6A, 6B, 6C). Overall, DC-AST VII (10 μM) alone activated dendritic cells, but the effect of DC-AST VII was diminished in the presence of LPS.

**FIGURE 6.**
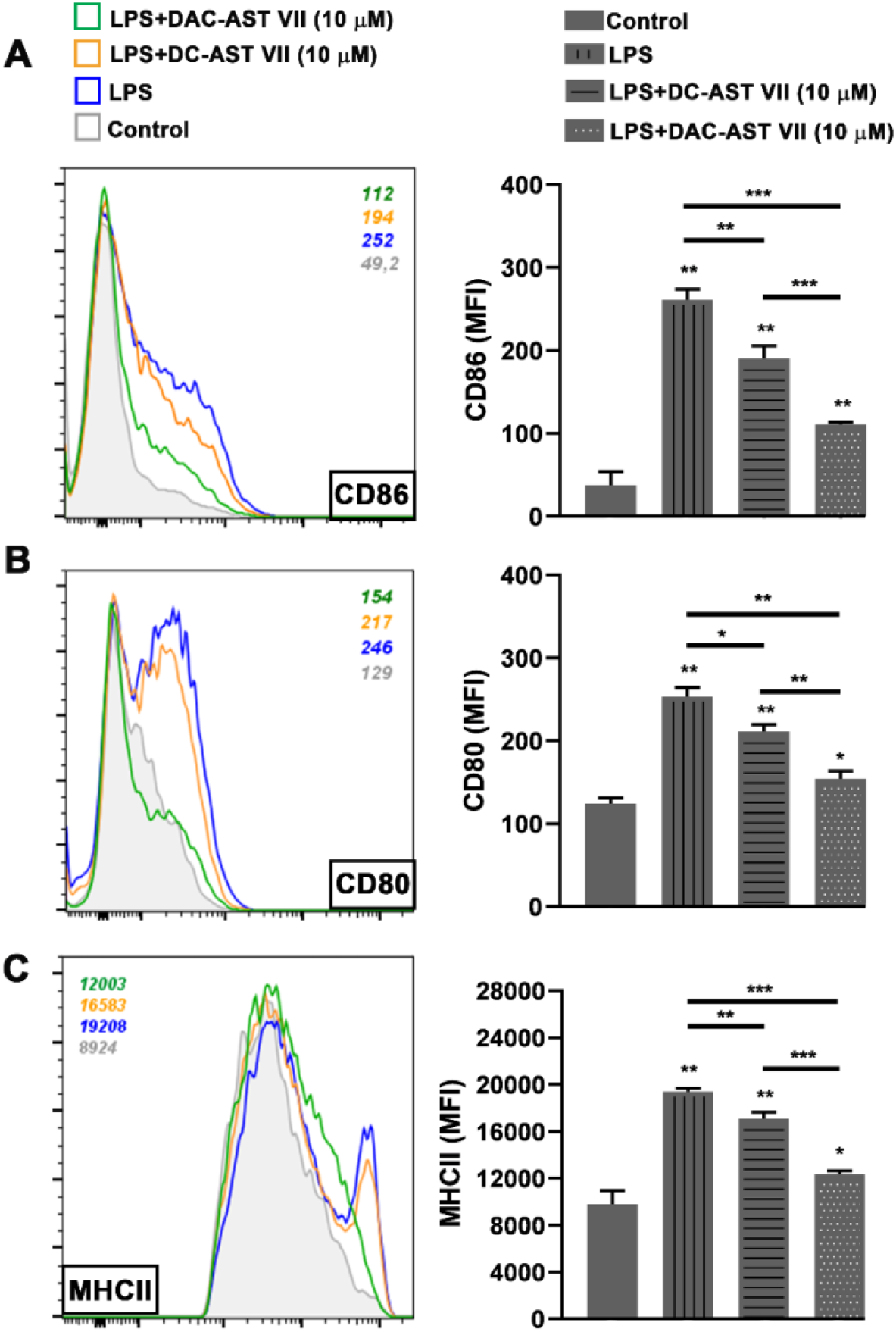
DC-AST VII and DAC-AST VII enhanced the maturation and activation of BMDCs in the presence of LPS. BMDCs generated from C57BL/6 mice bone marrow cells were treated with DC-AST VII or DAC-AST VII at the concentration of 10 μM for 24 h. The expression of **(A)** CD86, **(B)** CD80, and **(C)** MHC II on CD11c^+^ MHCII^+^ BMDCs was determined by flow cytometry. For A, B, and C, representative histograms (left panel) and summary graphs (right panel) are shown. Data shown are mean SD of triplicates and representative of two independent experiments with similar results. Statistically significant differences were performed between treated groups and control or LPS and, in treated groups between each other by One-way ANOVA and Student-t test. *p < 0.05, **p ≤ 0.01, ***p ≤ 0.001.

Next, we investigated whether the reduction in dendritic cell activation by the treatment of *Astragalus* saponins and LPS were correlated with their micelle formation properties. As saponins are amphiphilic compounds, due to the presence of a lipid-soluble aglycone and water-soluble sugar chain (30), they have the tendency to form micelles in aqueous solutions. Micelle formation is a concentration-dependent process that is characterized by a sharp transition at the critical micelle concentration (CMC). Below the CMC, the compounds are unassociated monomers or form a monolayer at the air-solvent interface, whereas above the CMC, the compounds assemble to form micelles (31) (Fig. 7A). To determine the micelle formation properties of *Astragalus* saponins, firstly, the compounds at various concentrations were mixed with fluorescence dye (dansyl chloride) overnight and relative fluorescence intensity at 508 nm were measured. Fluorescence intensity was plotted as a function of the logarithm of the concentrations of compounds, and each CMC value was calculated by the intersection of two linear lines. CMC values for AST VII, DC-AST VII and DAC-AST VII were calculated as 3.37 mM, 7.3 μM and 0.047 μM, respectively (Fig. 7B). Next, the particle size distribution of the compounds at the concentration above their CMC values was measured by dynamic light scattering (DLS). The hydrodynamic particle size for self-assembly micelles produced by AST VII (4000 μM) DC-AST VII (10 μM) and DAC-AST VII (10 μM) in aqueous solution was analyzed as 92 nm, 64 nm and 26 nm (Fig. 7C), respectively, and demonstrated a polydispersity index below than 0.5, attributed to a homogeneous size distribution. Moreover, zeta potential of these self-assembly micelles was higher than −20 mV, indicating the negatively charged, stable micelles were formed (Supplemental Table 3). These data showed that AST VII and its derivatives formed negatively charged self-assembly micelles in aqueous solution.

**FIGURE 7.**
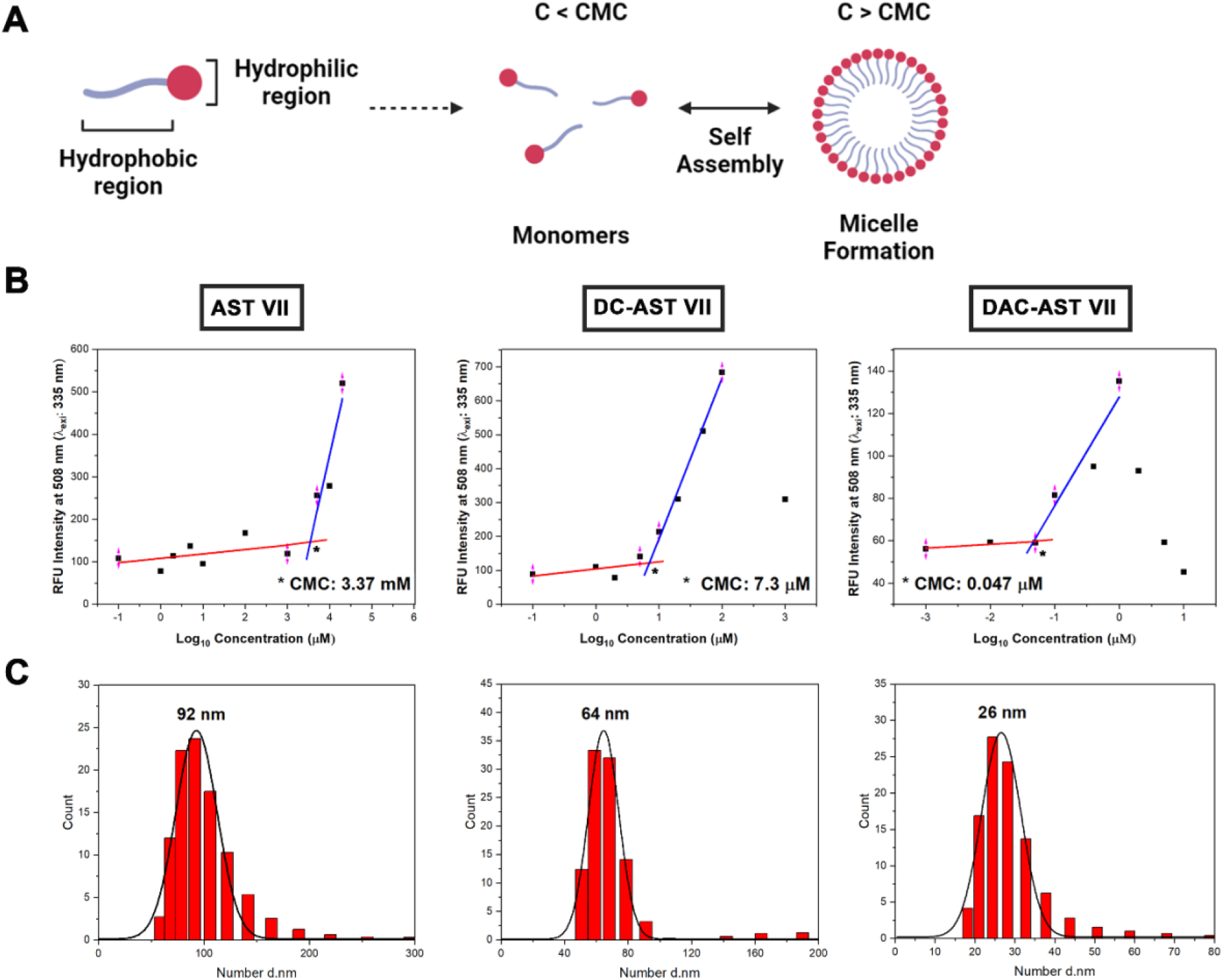
Self-assembling nanoparticles based on *Astragalus* saponins. (A) Schematic illustration of self-assembling micelle formation. (B) Relative fluorescence intensity at 508 nm vs logarithm of *Astragalus* saponins concentrations to determine critical micelle concentration (CMC). (C) Representative particle size distribution of self-assembling *Astragalus* saponins that were measured by dynamic light scattering (DLS) analysis. Data represent two or three independent experiments. *Created with* BioRender.com.

### AST VII and its derivatives activated CD4^+^ and CD8^+^ T cells in mixed leukocyte reaction (MLR)

To determine the impact of DC activation by AST VII and its derivatives on the ability of DCs to activate T cells, we performed a mixed leukocyte reaction (MLR) assay. MLR is a simple and effective *in vitro* model to investigate T cell activation and proliferation (32). BMDCs generated from the bone marrow cells of BALB/c mice were treated with stimulating agents such as LPS, AST VII, DC-AST VII, or DAC-AST VII for 24 h. The next day, naive CD4^+^ or CD8^+^ T cells isolated from the spleens of C57BL/6 mice were co-cultured with BMDCs for 3 days. The activation state of T cells was evaluated by measuring CD44 expression on T cells by flow cytometry as CD44 is expressed on activated T cells. In addition, CD44 enhances the stability of the DC-T cell interaction, the T cell proliferation to TCR signaling *in vitro*, and the cytokine response (33, 34).

Almost all tested compounds demonstrated an increase in the expression of CD44 on CD4^+^ and CD8^+^ T cells based on their structural features (Fig. 8A, 8B). In CD8^+^ T cells, LPS, LPS+AST VII (5 μM), and DAC-AST VII (10 μM) increased the expression of CD44 compared to the control (untreated) group (*p* < 0.05). DAC-AST VII was more potent in the activation of CD8^+^ T cells than LPS alone (*p* < 0.05). Interestingly, AST VII alone and DC-AST VII did not significantly augment CD44 on CD8^+^ T cells. Moreover, the levels of CD44 after LPS+AST VII, LPS alone or AST VII alone were comparable (n.s.)(Fig. 8A).

**FIGURE 8.**
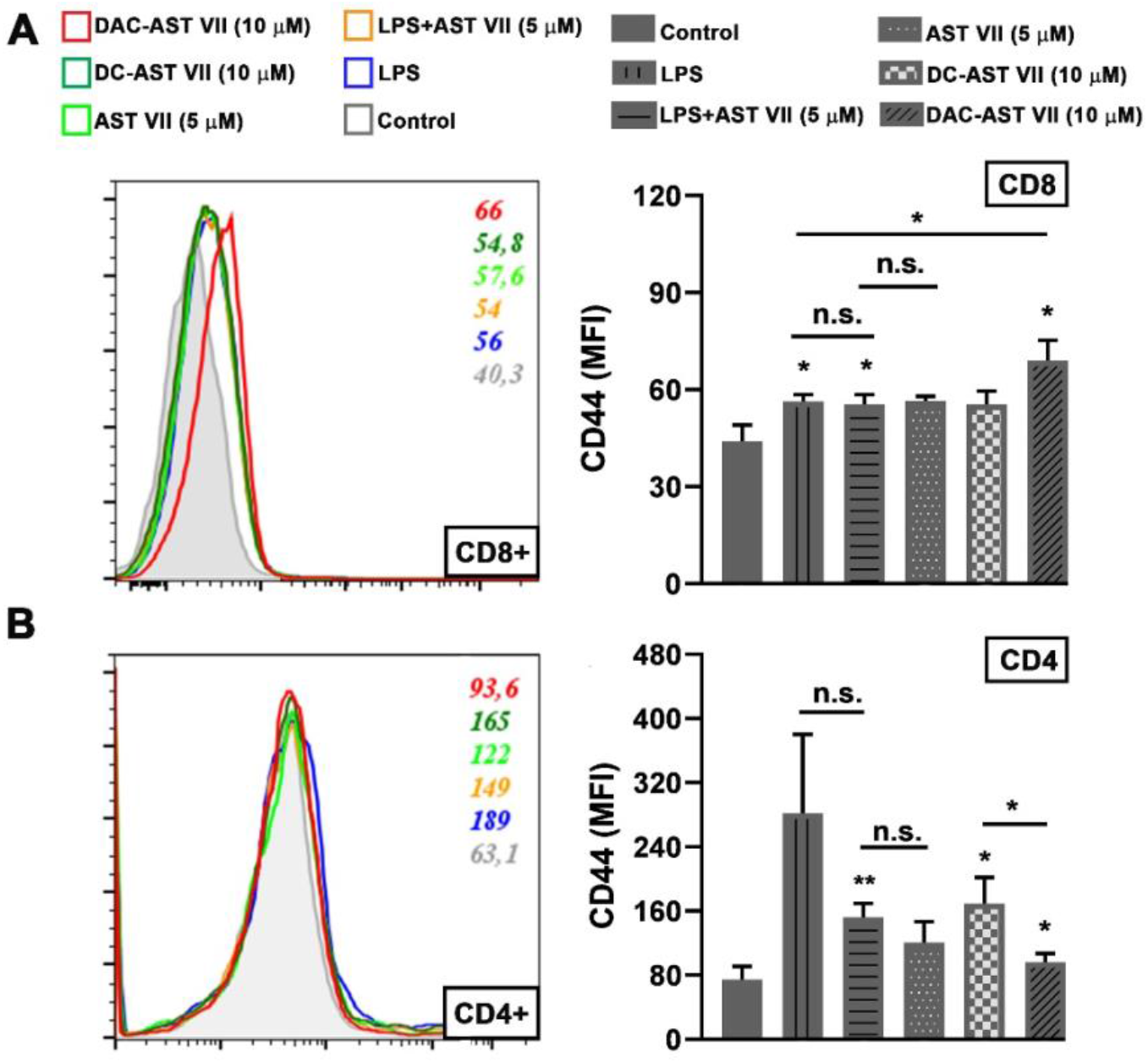
AST VII and its derivatives (DC-AST VII and DAC-AST VII) activated T cells in MLR. BMDCs derived from BALB/c mice were treated with LPS, AST VII, DC-AST VII, or DAC-AST VII for 24 h. The next day, BMDCs were co-cultured with naive CD4^+^ and CD8^+^ T cells isolated from the spleens of C57BL/6 mice for 3 days. The expression of CD44 on (**A**) CD8^+^ T cells and on (**B**) CD4^+^ T cells was determined by flow cytometry. For A and B, representative histograms (left panel) and summary graphs (right panel) are shown. Data shown are mean SD of triplicates and representative of three independent experiments with similar results. Statistically significant differences were analyzed between treated groups and control (untreated) and, in treated groups between each other by One-way ANOVA and Student-t test. *p < 0.05, **p ≤ 0.01, n.s. (not statistically significant).

In the case of CD4^+^ T cells, although LPS alone or AST VII alone did not have any statistically significant effect, AST VII and LPS co-treatment augmented CD44 expression compared to the control (untreated) group (*p* ≤ 0.01). As in the case of CD8^+^ T cells, there was no statistically significant differences in CD4^+^ T cell activation between neither LPS alone and LPS+AST VII nor AST VII alone and LPS+AST VII. In contrary, DC-AST VII (10 μM) and DAC-AST VII (10 μM) upregulate CD44 expression compared to the control group (*p* < 0.05). Moreover, DC-AST VII (10 μM) was more effective in CD4^+^ T activation compared to DAC- AST VII (10 μM) (*p* < 0.05). Our data demonstrate that the BMDCs treated with LPS+AST VII and DAC-AST VII activated both CD4^+^ and CD8^+^ T cells. DC-AST VII was only able to activate CD4^+^ T cells. Taken together, these data indicate that AST VII needs LPS to activate T cells, whereas both of the newly synthesized compounds have an ability to activate CD4^+^ T cells on their own.

## Discussion

Adjuvants are one of the essential components in modern vaccines that are capable of initiating innate immune response and subsequently adaptive immune response (35). The adjuvant potential of plant-derived compounds is widely investigated because of their abilities to enhance proper immune responses with limited toxicity (36). Saponins, a class of plant-derived adjuvants, trigger powerful immune response through Th1/Th2 balanced cytokine and antibody response (35). In the present study, the immunomodulatory activities of the purified *Astragalus* saponin, Astragaloside VII (AST VII), and its newly synthesized analogs DC-AST VII and DAC-AST VII were investigated *in vitro*.

DC-AST VII, having two glucuronic acid at C6 and C25 at the aglycone structure, represents a more hydrophilic structure than AST VII. On the other hand, DAC-AST VII with the dodecyl hydrocarbon chains extending from the glucuronic acid moieties at C6 and C25 on the aglycone forms the hydrophobic analog of AST VII. While DC-AST VII did not demonstrate cytotoxicity in cancerous cell lines as AST VII (16), DAC-AST VII showed cytotoxicity, indicating that lipophilicity is an important parameter for their cytotoxicity. Moreover, the compounds did not cause hemolysis on human red blood cells except DAC-AST VII (50 μM), which demonstrated slight hemolysis. The lysis of erythrocytes is one of the biological characteristics of saponins and it is a limiting factor for the utilization of these compounds in clinics (37). Therefore, our data indicates that AST VII and its derivatives have an advantage over other saponin compounds as hemolytic activity is one of the drawbacks in vaccine development.

To determine the cytokine release profiles of AST VII and its derivatives, Th1/Th2/Th17 cytokines were analyzed in hWB cells co-treated with PMA-ionomycin and the compounds. QS-21, a widely used saponin based adjuvant was also tested for comparison. In 1/10 diluted hWB, QS-21 and AST VII substantially increased IL-17A production; however, AST VII derivatives did not show such an effect except at particular concentrations. In 1/20 diluted hWB, IL-17A was not detected in the cell culture supernatant following AST VII treatment, possibly resulting from the consumption of IL-17A via IL-17 receptor. This consumption by specific receptor *in vitro* has been also suggested as the possible reason for undetectable IL-4 in cell culture supernatant (38, 39).

A contradicting result to our previous studies (16, 17, 40) was obtained with Th1 cytokines (IL-2 and IFN-γ), as the saponin treatment unexpectedly suppressed these cytokines in contrast to P+I treatment in 1/20 diluted hWB. As AST VII was shown to be a promising compound for IL-2 production (40), it was also more effective in IL-2 production in this study compared to its derivatives and QS-21. Although some triterpenoid saponins, like Ginsenoside Rg1 and Ginsenoside Rd, were shown to induce Th2 cytokines, AST VII and its derivatives did not stimulate IL-4 production, in parallel with previous reports (16, 17). Therefore, we conclude that AST VII and its derivatives do not lead to a Th2 immune response. Vitoriano-Souza et al. investigated the cytokine patterns of saponin adjuvant (SAP) in sensitized mouse skin at 12, 48 and 168 hours. At 12 h, the production of TNF-α, IL-6, IL-2, IL-17, and IL-4 cytokines were induced. At 48 h, while TNF-α, IFN-γ and IL-2 levels were increased, IL-6, IL-17 and IL-4 levels were decreased. At 168 h, all cytokine levels were comparable to the baseline value (41). Therefore, the treatment time is also crucial to observe the production of the desired cytokines.

Another prominently induced cytokine by AST VII and derivatives was IL-1β. In 1/10 diluted hWB in the presence of PMA-ionomycin, all compounds enhanced IL-1β production in a dose-dependent manner. However, only DAC-AST VII increased IL-1β production in 1/20 diluted hWB in the presence of PMA-ionomycin. The difference between AST VII and DAC- AST VII in terms of IL-1β inducing in hWB cells could be due to a differing activation capacity of these compounds on distinct immune cells as hWB contains various immune cells (24). In addition, one of the reasons for differences in IL-1β induction by saponin compounds could be a result of their hydrophobic property. Calculated partition coefficients (clogP) clearly demonstrate a lipophilic order of test compounds [DAC-AST VII (clogP: 11.41) > AST VII (clogP: 1.29) > DC-AST VII (clogP: 0.34)], which imply that polarity is indeed an important factor for IL-1β induction in hWB cells. IL-1β helps naive T cells to differentiate into Th17, promotes IL-17 producing memory CD4^+^ T cells and is a profound inducer of IL-17^+^ T cell differentiation along with TGF-β(42, 43). Nalbantsoy et al. reported that AST VII in the presence of LPS was able to stimulate TGF-β production (17). Based on these results, it is possible that AST VII promotes IL-17A production in the presence of IL-1β and TGF-β, whereas analogs of AST-VII are not able to exhibit such an effect on IL-17A secretion except for DC-AST VII and DAC-AST VII at 2 and 16 μg/mL concentrations, respectively.

IL-1β production and its secretion progress in three steps: a) priming: production of pro-IL-1β; b) processing of pro-IL-1β to mature IL-1β; and c) release of IL-1β from the cells (44). The prominent induction of IL-1β lead us to investigate the role of these compounds in innate cell activation. As the priming step requires stimulation of the TLR/NF-κB pathway, and as AST VII alone cannot activate NF-κB (45), LPS was used to activate pro-IL-1β in BMDCs and BMDMs (46). Co-stimulation of BMDCs/BMDMs with LPS and AST VII or its analogs lead to a higher production of IL-1β compared to LPS alone, and the most potent compound was AST VII. However, low levels of IL-1β was induced by DAC-AST VII treatment in BMDMs, suggesting possible cytotoxicity of DAC-AST VII on BMDMs. Astragaloside IV (AST IV), another cycloartane type triterpene glycoside containing xylose and glucose moieties at C-3 and C-6 on the aglycone backbone, has anti-inflammatory properties. Li et al. demonstrated that AST IV inhibited the release of the pro-inflammatory cytokines TNF-α and IL-1β and the activation of NF-κB in LPS induced epithelial cells *in vitro* and *in vivo*. Moreover, treatment of AST IV alone did not alter the gene expressions of pro-inflammatory cytokines and adhesion molecules (47, 48). In addition, AST IV suppressed the RhoA/NLPR3 signalling pathway in sepsis-induced mice (49). According to these reports, the presence of glucose at C-25 on the triterpenoid aglycone of AST VII, possibly changes the ability to induce IL-1β production, signifying the importance of the tridesmosidic nature of AST VII. QS-21 (2 μg/mL) in the presence of the TLR4 agonist MPLA was reported to enhance caspase-1/11 and NLPR3 dependent IL-1β production. QS-21 induced IL-1β titers were higher than alum, a well-known inflammasome activator (11). Moreover, Roix et al. investigated whether inflammasome activation was a general phenomenon for saponins or not. Macrophages were treated with different saponins such as QS-21, digoxin, sapindoside A, hedaracoside C or β-escin in combination with MPLA. Only QS-21, which is isolated from *Quillaja saponaria*, elicited an IL-1β response, indicating that inflammasome activation could be specific to *Quillaja saponins*. To test this hypothesis, *Quillaja* saponin (Quil-A®) and VET-SAP® were administrated to BMDCs/ BMDMs, and both showed NLRP-3 dependent IL-1β secretion (11). Therefore, AST VII and derivatives may be inducing IL-1β responses in BMDCs through the activation of the inflammasome. Further studies are needed to test this hypothesis.

IL-1β can act as a maturation factor for DCs in terms of IL-12 production and expression of CD86, MHC I, and ICAM-1 (50). Thus, we carried out experiments to evaluate the activation status of dendritic cells following treatment with AST VII and its derivatives. Expression of the activation markers CD80, CD86, and MHC II and the secretion of IL-12 were investigated in BMDCs. AST VII alone did not induce the activation of dendritic cells, either BMDCs or in splenocytes, in terms of co-stimulatory molecule expression and IL-12 secretion *in vitro*. In the presence of LPS, AST VII was capable to induce CD86, CD80, and MHC II upregulation and and IL-12 production comparable to LPS alone. This suggests that AST VII is not effective by itself to elicit immunostimulatory activity *in vitro*, and requires a co-stimulatory agent such as LPS. A previous study carried out by Marty-Roix et al. showed that QS-21 alone did not lead to the activation of murine bone marrow-derived dendritic cells and macrophages directly (11). QS-21 in liposomal formulation with MPL, another TLR-4 agonist, augmented MHC II and CD86 expression on human monocyte derived dendritic cells (10).Therefore, our study shows that AST VII influence on immune responses is comparable to QS-21. On the other hand, DC- AST VII alone increased the expression of CD86, CD80, and MHC II on BMDCs, whereas DAC-AST VII alone had no effect on the expression of these markers. The upregulation of CD86, CD80, and MHC II induced by DC-AST VII was higher than by DAC-AST VII. This suggests that carboxylic acid moieties on the structure or more hydrophilic character of DC- AST VII may have positive effects on the dendritic cell activation. When LPS was added to the culture along with DC-AST VII and DAC-AST VII, the expression of CD80, CD86, and MHC II were significantly reduced compared to LPS alone. Based on CMC values, it can be concluded that AST VII is mainly a monomer, whereas DC-AST VII and DAC-AST VII form self-assembly micelles at the treatment concentrations. Moreover, LPS has a very low critical micelle concentration of 10^−9^ mol (51). Only its monomers can bind to TLR4/MD2 heterodimer, and further transduce the signal (52). Thus, AST VII may be facilitating the transfer and binding of LPS to TLR4/MD2. In the case of AST VII derivatives, micelle formation can lead to the encapsulation of LPS as hydrophobic properties of LPS enables itself to be entrapped into micelles easily. The difficulties in the release of LPS from micelles led to the reduction of dendritic cell activation. As micelles derived from DAC-AST VII contains the dodecyl acyl chains in its inner core, the stability of LPS entrapped in the micelle will be increased, causing a stronger reduction of the dendritic cell activation compared to DC-AST VII.

Considering the IL-1β secretion and up-regulation of MHC II and co-stimulatory molecules following the co-treatment with LPS, a synergistic effect of AST VII on TLR/NLR interaction might be the reason for the augmentation of the innate immune response. Further, AST VII may have a role in the induction of NLRs through the interaction with the cell membrane. We postulate that triterpenic AST VII can rearrange the cell membrane organization by intercalating its aglycone backbone in the phospholipid bilayer and extend sugar moieties to the extracellular space. This alteration in the cell membrane can be sensed by dendritic cells as a pathological condition and transduce the signal for its maturation and activation. Moreover, such an effect can activate Syk kinases to assemble the inflammasome and promotes secretion of IL-1β. Since QS-21 has been shown to activate Syk in monocyte derived dendritic cells (10), it is plausible that AST VII may be inducing IL-1β production through Syk signaling as well.

A few adjuvants are known to induce effective cellular immune response (53). As CD44 is one of the indicators of T cell activation, the impact of AST VII and its derivatives on CD44 upregulation was analyzed. LPS + AST VII or DAC-AST VII enhanced CD44 expression on both CD4^+^ and CD8^+^ T cells. DAC-AST VII, the lipophilic analog of AST VII because of dodecyl conjugation, was better at CD8^+^ T cell activation while DC-AST VII was more potent in CD4^+^ T cell activation. Overall, our results indicate that AST VII needs LPS to activate T cells, but AST VII analogs can activate T cells without LPS co-treatment, suggesting different action mechanisms for AST VII and derivatives. Imine-formation between carbonyl group on DC-AST VII and DAC-AST VII and amino groups, could be have a role in the activation of T cells. This imine or Shiff-base formation by reacting carbonyl groups with amino groups, likely from CD2 receptor on T cells, delivers a signal that replaces the one derived from the interaction between CD80/86 ligands and CD28 receptor on dendritic cells and T cells, respectively (54).

In a previous study, BMDCs were co-treated with OVA antigen and Matrix C, which is an adjuvant based on ISCOM that contains *Quillaja* saponin, phospholipids, and cholesterol and later co-cultured with OVA-specific CD8+ T cells (OT-I cells). Matrix C upregulated CD44 expression on OT-I cells compared to OVA treatment without the adjuvant (55). Marciani et al. stated that the lipophilic acyl side chain of *Quillaja* saponin was an important structural feature for stimulation of cytotoxic T cell (CTL) proliferation (56). Moreover, it was reported that dodecylamine conjugated QS-21, namely GPI-0100, induced Th1 and CTL responses. The lipophilic character of GPI-0100 was proposed to enable the compound to open a transient pore for delivery of exogenous proteins to the cytosol and further enhance MHC I restricted CTL responses (7). In another study, the QS-21 derivative 3, prepared by modification of the glucuronic acid carboxyl with ethylamine, was shown to enhance CTL responses comparable to QS-21 in mitomycin-C treated OVA-specific E.G7-OVA cells (57). On the other hand, the QS-21 derivatives 2 and 4, prepared by the modification of glucuronic acid carboxyl with glycine and ethylenediamine, was shown to increase CTL response but not as high as QS-21. Increases in the lipophilicity of the compound influenced CTL responses (57). In this study, in a similar manner, we introduced the dodecylamine onto glucuronic acid residues and increased the lipophilicity of AST VII and demonstrated that the CTL (CD8^+^ T cell) response was improved. This may be due to the enhanced translocation of the antigen into the cytosol, leading to more efficient antigen presentation by MHC I molecules to CD8^+^ T cells, which in turn results in TCR:MHC I engagement and upregulation of CD44. As CD44 expression can be regulated by pro-inflammatory cytokines such as IL-1β (58), the IL-1β secretion from dendritic cells might be the cause for the CD44 upregulation on T cells. Therefore, further studies are warranted to show the effects of IL-1β on CD44 gene regulation.

In conclusion, our study demonstrates that cycloartane type saponins, AST VII and the newly synthesized derivatives DC-AST VII and DAC-AST VII, induce IL-1β production in hWB cells and BMDCs. Robust IL-1β production by these compounds was associated with the maturation and activation of dendritic cells, CD4^+^ and CD8^+^ T cells. Modification of the compounds towards more or less polar derivatives altered the activation status of CD4^+^ and CD8^+^ T cell responses. As there is a great need for adjuvants that can effectively enhance cellular immune response, the compounds presented here could be good alternatives to other saponin-based adjuvants. This study is a first step to evaluate the mechanism of action of AST VII and subsequent studies towards a rational design of AST VII based adjuvant analogs/vaccine formulations to be utilized in prophylactic and therapeutic vaccines are in progress.

## Acknowledgement

We thank Sinem Günalp for excellent technical assistance and Sinem Yılmaz for helping with cell viability experiments. We also thank Dr. Gerhard Wingender for valuable scientific contributions.

