## Supplementary Material for "*Astragalus* Saponins, Astragaloside VII and Newly Synthesized Derivatives, Induce Dendritic Cell Maturation and T Cell Activation Through IL-1β Production"

**Supplemental Table 1.** Semi-synthesis of AST VII Analogs (DC-AST VII and DAC-AST VII) and structure elucidation by NMR and MS

| Compound | Semi-synthesis protocol | HR-ESI-MS m/z | $^{1}$H NMR (400 MHz) | $^{13}$C NMR (100 MHz) |
|---|---|---|---|---|
| DC-AST VII | AST VII (1000 mg, 1.06 mmol, 1 equiv.), NaBr (109 mg, 1.06 mmol, 1 equiv.) and TEMPO (40 mg, 0.212 mmol, 0.2 equiv.) were dissolved in distilled water (pH 11 adjusted with 1 N NaOH). NaOCl (6.3 mL, 4.66 mmol, 4.4 equiv) was slowly added to the reaction mixture and stirred at 0°C. After 4 h, the reaction was quenched by addition of distilled water and neutralized by 1 M HCl. Reaction mixture was extracted (x3) with *n*-butanol and water phase was evaporated at 50°C in rotary evaporator. Further purification was done by VLC (vacuum liquid chromatography) loaded with reversed-phase silica gel (RP-18, 25 g) using MeOH/$H_2O$ gradient (15:85, 20:80, 35:65, 40:60; 50:50, 55:45, 100:0). | 995.38897 ([M+Na-2H]$^-$) $C_{47}H_{72}O_{21}Na$= 995.444. | **$^{1}$H-NMR ($D_2O$)** δ 4.72 (1H, dd, *J= 6, 4.5 Hz*, H-16), 4.70 (1 H, d, *J= 8.1 Hz*, H-1’’’), 4.56 (1 H, d, *J= 7.9 Hz*, H-1’’), 4.50 (1 H, d, *J= 7.8 Hz*, H-1’), 3.97 (1 H, dd, *J= 15.5, 6.4 Hz*, H-24), 3.96 (1 H, d, *J= 5.8 Hz*, H-5’), 3.73 (2 H, s, H-5’’, H-5’’’), 3.69 (1 H, d, *J= 5.3 Hz*, H-6), 3.68 (1 H, d, *J= 9.1 Hz*, H-4’), 3.56 (1 H, d, *J= 4.1 Hz*, H-4’’), 3.54 (1 H, d, *J= 2.7 Hz*, H-3’), 3.52 (1 H, d, *J= 2.9 Hz*, H-3’’), 3.50 (1 H, s, H-3’’’), 3.39 (2 H, d, *J= 4.4 Hz*, H-2’, H-3), 3.33 (1 H, s, H-5’), 3.31 (1 H, d, *J= 6.2 Hz*, H-2’’), 3.30 (1 H, d, *J= 8.5 Hz*, H-2’’’), 3.56 (1 H, d, *J= 4.1 Hz*, H-4’’’), 2.46 (2 H, d, *J= 8 Hz*, H-17, H-22), 2.16 (1 H, s, H-23), 2.12 (2 H, d, *J= 9.2 Hz*, H-23, H-11), 2.08 (1 H, m, H-15), 1.97 (1 H, d, *J= 7.7 Hz*, H-2), 1.95 (1 H, d, *J= 7.7 Hz*, H-7), 1.84 (1 H, d, *J= 12.2 Hz*, H-8), 1.75 (1 H, d, *J= 12.5 Hz*, H-22), 1.74 (1 H, d, *J= 12.5 Hz*, H-12), 1.68 (1 H, s, H-5), 1.67 (2 H, s, H-1, H-2), 1.64 (1 H, s, H-12), 1.45 (1 H, d, *J= 10.1 Hz*, H-7), 1.44 (1 H, d, *J= 10.1 Hz*, H-15), 1.39 (3 H, s, H-26), 1.32 (3 H, s, H-28), 1.30 (3 H, s, H-27), 1.29 (6 H, s, H-21, H-18), 1.22 (1 H, s, H-1), 1.16 (1 H, s, H-11), 1.06 (3 H, s, H-29), 1.01 (3 H, s, H-30), 0.7 (1 H, s, H-19), 0.4 (1 H, s, H-19). | **$^{13}$C-NMR ($D_2O$)** δ 176.1 (s, C-6’’), 175.7 (s, C-6’’’), 105.5 (d, C-1’), 102.5 (d, C-1’’), 96.8 (d, C-1’’’), 89.3 (d, C-3), 87.9 (s, C-20), 81.6 (d, C-24), 80.3 (d, C-6), 79.7 (s, C-25), 76.6 (d, C-5’’), 76.3 (d, C-5’’’), 76.1 (d, C-3’), 75.9 (d, C-3’’, C-3’’’), 73.8 (d, C-16), 73.7 (d, C-2’’’), 73.6 (d, C-2’), 73.1 (d, C-2’’), 71.8 (d, C-4’’, C-4’’’), 69.4 (d, C-4’), 65.1 (t, C-5’), 57.0 (d, C-17), 51.6 (d, C-5), 46.3 (d, C-8), 45.6 (s, C-13), 44.8 (s, C-14), 44.2 (t, C-15), 41.5 (s, C-4), 34.6 (t, C-7), 34.3 (t, C-22), 32.9 (t, C-12), 31.9 (t, C-1), ), 30.7 (t, C-19), 29.2 (t, C-2), 29.1 (s, C-10), 27.4 (q, C-28), 27.3 (q, C-21), 25.7 (t, C-11), 25.6 (t, C-23), 21.2 (q, C-18), 20.4 (s, C-9), 24.7 (q, C-26), 21.5 (q, C-27), 19.3 (q, C-30), 15.7 (q, C-29). |

| | | | | |
|---|---|---|---|---|
| DAC-AST VII | Dicarboxylic AST-VII (50 mg, 0.0513 mmol, 1 equivalent) was dissolved in pyridine. DIPEA (27 mg, 0.2052 mmol, 4 equiv.), HOBt (16 mg, 0.1026 mmol, 2 equiv.), EDC (30 mg, 0.1539 mmol, 3 equiv.) were added and the reaction mixture was stirred for 1 h at room temperature. 1 h later, free dodecylamine (23 mg, 0.1231 mmol, 2.4 equiv.) was added dropwise and reaction mixture was heated to 60°C. After 6 h, the reaction was quenched by addition of distilled water and extracted (x3) with ethyl acetate. Ethyl acetate fraction was evaporated at 50°C in rotary evaporator. DAC-AST VII was purified by silica gel open-column chromatography (30 g) eluting with $CHCl_3$: MeOH: $H_2O$ (80:20:2). | 1331.89196 ($[M+Na]^+$) $C_{71}H_{124}O_{19}Na$ =1331.88. | **$^1$H-NMR (400 MHz, d5)** δ 8.13 (1H, t, *J = 6.2 Hz*, H-1$^{v}$), 8.09 (1H, t, *J = 6.1 Hz*, H-1$^{iv}$), 5.12 (1H, d, *J = 7.8 Hz*, H-1’’’), 4.99 (1H, s, H-16), 4.95 (1H, d, *J = 7.8 Hz*, H-1’’), 4.86 (1H, d, *J =7.4 Hz*, H-1’), 4.36 (1H, dd, *J = 11.0, 4.7 Hz*, H-5’), 4.35 (1H, s, H-5’’), 4.35 (1H, d, *J = 5.3 Hz*, H-5’’’), 4.26 (2H, dd, *J = 7.8, 4.7 Hz*, H-3’’, H-4’’), 4.26 (1H, m, H-2’’’), 4.22 (1H, m, H-4’), 4.21 (2H, d, *J = 4.9 Hz*, H-3’’’, H-4’’’), 4.15 (1H, t, H-3’), 4.04 (2H, m, H-2’, H-2’’), 3.95 (1H, dd, *J = 8.5, 6.1 Hz*, H-24), 3.83 (1H, m, H-6), 3.72 (1 H, d, *J = 10.7 Hz*, H-5’), 3.64 (2H, m, H-2$^{v}$), 3.53 (1H, dd, *J = 11.8, 4.4 Hz*, H-3), 3.46 (2H, m, H-2$^{iv}$), 2.76 (1H, t, *J = 9.9 Hz*, H-22),2.53 (1H, m, H-17), 2.39 (1H, d, *J = 12.4 Hz*, H-11), 2.34 (1H, s, H-23), 2.26 (1H, dd, *J = 12.3, 7.9 Hz*, H-15),2.15 (1H, dd, *J = 8.5, 3.5 Hz*, H-7), 2.01 (1H, s, H-8), 1.99 (3H, s, H-28), 1.95 (1H, m, H-23), 1.95 (1H, s, H-11), 1.88 (1H, d, *J = 8.8, 4.4 Hz*, H-5), 1.84 (1H, d, *J = 10.1*, H-7), 1.80 (1H, m, H-1), 1.78 (1H, d, *J = 6.4 Hz*, H-15), 1.74 (1H, d, *J = 7.3 Hz*, H-12), 1.73 (1H, m, H-2), 1.65 (1 H, s, H-22), 1.64 (3H, s, H-26), 1.62-1.82 (36H, m, H-3$^{iv}$-11$^{iv}$, H-3$^{v}$-11$^{v}$), 1.56 (2H, d, *J = 12.1 Hz*, H-1, H-12), 1.44 (3H, s, H-27), 1.42 (3H, s, H-18), 1.37 (3H, s, H-29), 1.32 (3H, s, H-21), 1.29 (4H, m, H-12$^{iv}$, H-12$^{v}$), 1.26 (1H, m, H-2), 1.12 (3H, s, H-30), 0.88 (6H, s, H-13$^{iv}$, H-13$^{v}$), 0.61 (1H, d, *J = 3.9 Hz*, H-19), 0.25 (1H, d, *J = 4.1 Hz*, H-19). | **$^{13}$C-NMR (100 MHz, d5)** δ 171.5 (s, C-6’’), 171.4 (s, C-6’’’), 108.2 (d, C-1’), 105.2 (d, C-1’’), 99.1 (d, C-1’’’), 88.9 (d, C-3), 87.7 (s, C-20), 82.5 (d, C-24), 79.7 (s, C-25), 79.5 (d, C-6), 79 (d, C-3’), 78.9 (d, C-3’’, C-2’’’), 78.3 (d, C-3’’’), 76.6 (d, C-5’’), 76.4 (d, C-5’’’), 75.4 (d, C-2’’), 74.9 (d, C-2’), 74.4 (d, C-4’’’), 74.2 (d, C-4’’), 74.1 (d, C-16), 71.8 (d, C-4’), 67.6 (t, C-5’), 58.6 (d, C-17), 52.9 (d, C-5), 46.8 (s, C-13), 46.7 (t, C-15), 45.9 (d, C-8), 45.9 (s, C-14), 43.2 (s, C-4), 39.8 (t, C-2$^{v}$), 35.7 (t, C-22), 34.7 (t, C-7), 33.9 (t, C-12), 32.7 (t, C-1,C-12$^{iv}$,C-12$^{v}$), 30.2 (t, C-2), 30-31 (t, C-3$^{v}$- C-11$^{v}$), 29.5 (d, C-10), 28.9 (t, C-19), 28.2 (q, C-28), 27.8 (q, C-21), 27.8-31 (t, C-3$^{iv}$- C11$^{iv}$), 26.7 (t, C-11), 26.5 (t, C-23), 25.4 (q, C-26), 23.5 (q, C-27), 21.6 (q, C-18), 21.5 (s, C-9), 20.6 (q, C-30), 17.1 (q, C-29), 14.6 (q, C-13$^{iv}$, C-13$^{v}$). |

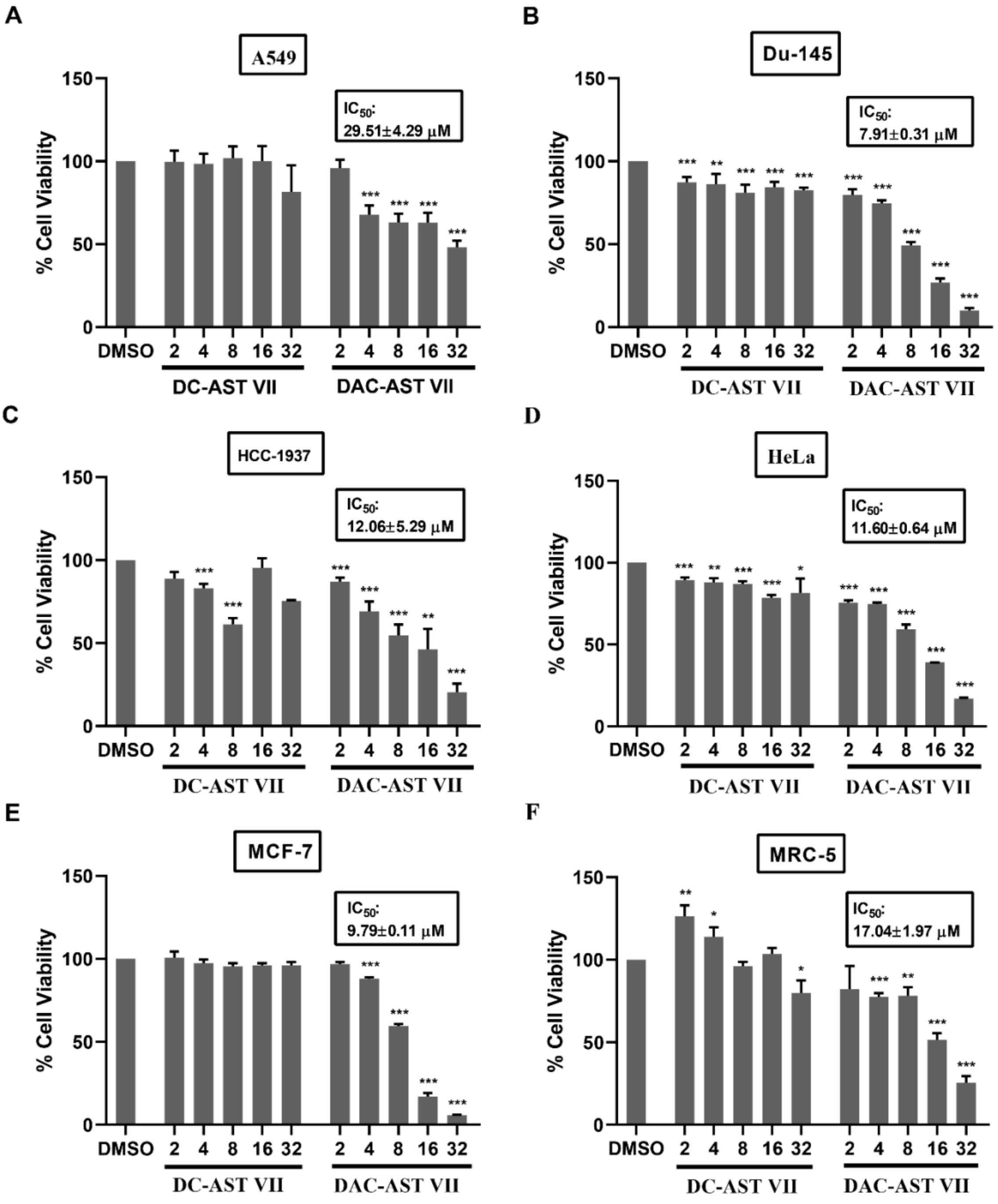

**Supplemental Figure 1.** Determination of cell viability and $IC_{50}$ values in cancer and healthy cell lines treated with DC-AST VII and DAC-AST VII. The cell viability (%) and related $IC_{50}$ values in different cancer **(A)** A549, **(B)** Du145, **(C)** HCC-1937, **(D)** HeLa, **(E)** MCF-7 and healthy **(F)** MRC-5 cell lines treated with DC-AST VII and DAC-AST VII were represented. A549, Du145, HCC-1937, HeLa, MCF-7 and MRC-5 cell lines were treated with DC-AST VII and DAC-AST VII at the concentrations of 2 to 32 µM for 48 h. The cell viability was analyzed by MTT assay and calculated compared to DMSO. DMSO alone was used as vehicle control, with the results representing average values from individual experiment, each performed in triplicate. Statistical analyses were performed between treated groups and DMSO control using One-way ANOVA and Dunnett's multiple comparisons test. $*p < 0.05$, $**p < 0.01$, $***p < 0.001$.

**Supplemental Table 2.** Hemolytic activity of DAC-AST VII and DC-AST VII. Statistically significant differences of treated groups versus saline control are indicated. *$p < 0.05$, **$p < 0.01$, ***$p < 0.001$ by one-way ANOVA and multiple comparion test.

| Groups | Hemolysis (%) |
|---|---|
| Saline | 0 |
| Distilled Water | 100*** |
| DAC-AST VII (μM) | |
| 250 | 1,377 ± 0,031 |
| 50 | 2,411 ± 1,922** |
| 10 | 1,436 ± 0,658 |
| 2 | 0,762 ± 0,053 |
| 0.4 | 1,199 ± 0,163 |
| DC-AST VII (μM) | |
| 250 | 1,246 ± 0,553 |
| 50 | 0,763 ± 0,012 |
| 10 | 1,616 ± 0,897* |
| 2 | 0.755 ± 0,204 |
| 0.4 | 1,413 ± 0,396 |

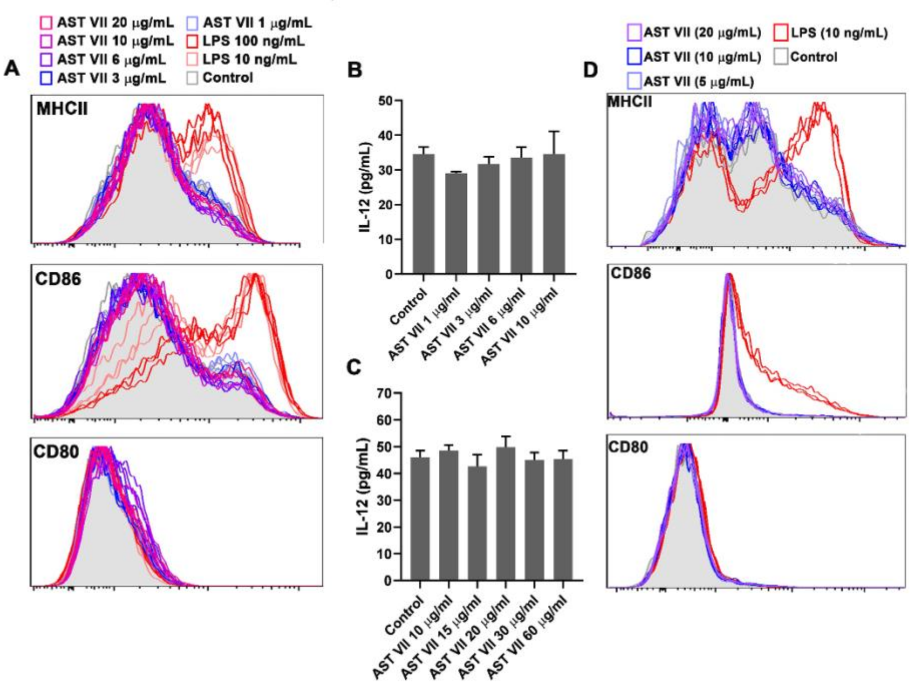

**Supplemental Figure 2.** AST VII alone did not induce the maturation and activation of BMDCs and splenic dendritic cells. BMDCs or splenocytes were treated with AST VII at the concentrations of 1 to 20 µg/mL for 24 h. The surface marker expression (**A** and **D**) by BMDCs and CD11c$^{+}$MHCII$^{+}$ dendritic cells in splenocytes were analyzed by flow cytometry. (**B** and **C**) IL-12 titers in the cell culture supernatant of BMDCs were measured by ELISA. Representative data from one of the two independent experiments, each performed in triplicate, are shown.

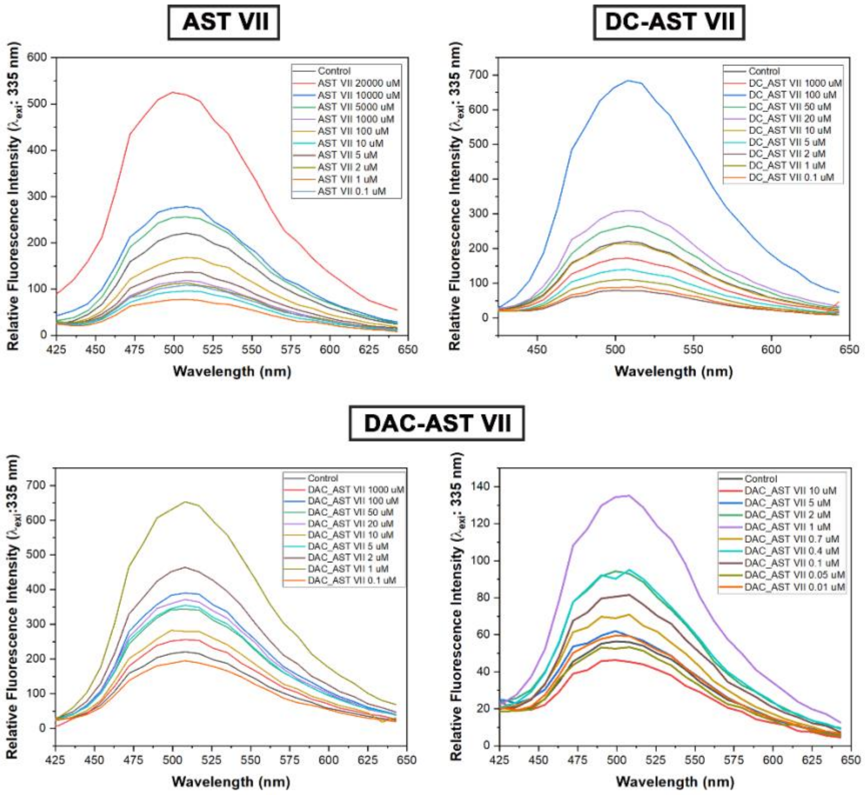

**Supplemental Figure 3.** Dansyl chloride shows increasing fluorescence intensity with increasing concentrations of AST VII and derivatives ($\lambda_{exitation}$: 335 nm).

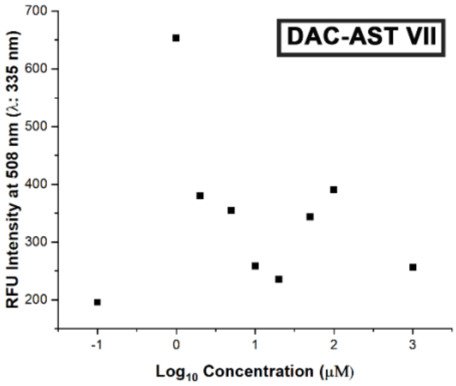

**Supplemental Figure 4.** Self-assembling nanoparticles based on DAC-AST VII. Relative fluorescence intensity at 508 nm vs logarithm of DAC-AST VII (0.1 to 1000 μM) to determine critical micelle concentration (CMC).

**Supplemental Table 3.** Zeta potential of AST VII and derivatives.

| Compounds | Zeta Potential (mV) |
|---|---|
| AST VII | -19.5 mV |
| DC-AST VII | -26.5 mV |
| DAC-AST VII | -23.4 mV |
